# Pyocin S5 import into *Pseudomonas aeruginosa* reveals a generic mode of bacteriocin transport

**DOI:** 10.1101/856047

**Authors:** Hannah M. Behrens, Edward D. Lowe, Joseph Gault, Nicholas G. Housden, Renata Kaminska, T. Moritz Weber, Catriona M A Thompson, Gaëtan L. A. Mislin, Isabelle J. Schalk, Daniel Walker, Carol V. Robinson, Colin Kleanthous

**Author notes:** Address for correspondence: Prof Colin Kleanthous, Department of Biochemistry, University of Oxford, South Parks Road, Oxford OX1 3QU, UK.

## Abstract

Pyocin S5 (PyoS5) is a potent protein bacteriocin that eradicates the human pathogen *P. aeruginosa* in animal infection models, but its import mechanism is poorly understood. Here, using crystallography, biophysical and biochemical analysis and live-cell imaging, we define the entry process of PyoS5 and reveal links to the transport mechanisms of other bacteriocins. In addition to its C-terminal pore-forming domain, elongated PyoS5 comprises two novel tandemly repeated kinked three helix bundle domains that structure-based alignments identify as key import domains in other pyocins. The central domain binds the lipid-bound common polysaccharide antigen, allowing the pyocin to accumulate on the cell surface. The N-terminal domain binds the ferric pyochelin transporter FptA while its associated disordered region binds the inner membrane protein TonB1, which together drive import of the bacteriocin across the outer membrane. Finally, we identify the minimal requirements for sensitizing *Escherichia coli* towards PyoS5, as well as other pyocins, and suggest that a generic pathway likely underpins the import of all TonB-dependent bacteriocins across the outer membrane of Gram-negative bacteria.

## Introduction

Bacteria living within communities do so through cooperation and antagonism. Forms of antagonism involving one bacterium killing another are important for maintaining the stable co-existence of bacteria within microbiomes deployed by pathogens and commensals alike to kill competitors (Granato *et al*, 2019). Antagonism occurs via several routes, the most common being bacteriocins, contact-dependent inhibition or type VI secretion. Of these, only the release of bacteriocins does not rely on physical contact between bacterial cells. Bacteriocin production generally occurs following a stress signal, such as DNA damage, inducing expression and release of the bacteriocin from auto-lysed cells (Kleanthous, 2010). The bacteriocin then diffuses through the medium to kill a neighbouring cell. Bacteriocins range in size, from small peptides to large proteins with both types currently being evaluated/developed as antimicrobials against multidrug resistant bacteria (Rios *et al*, 2016; Behrens *et al*, 2017). In many instances, however, developments are hindered by a lack of understanding as to how these molecules work. In the case of protein bacteriocins, extensive sequence diversification and homologous recombination further hamper efforts to find generic mechanisms of uptake. Here, we focus on the uptake mechanism of PyoS5, a protein bacteriocin that specifically targets the opportunistic human pathogen *P. aeruginosa* and shown recently in animal models to be more effective at clearing lung infections than tobramycin, the antibiotic generally used to treat *P. aeruginosa* in cystic fibrosis patients (McCaughey *et al*, 2016b). Through a structure-led approach, we deconstruct the energised uptake pathway of PyoS5 and show that its transport across the outer membrane likely represents the default pathway for all TonB-dependent bacteriocins.

There is a pressing need for new antibiotics against Gram-negative bacteria but in particular *P. aeruginosa* which has been designated a priority pathogen (WHO, 2017). The intrinsic low permeability of its outer membrane renders *P. aeruginosa* insensitive to many classes of antibiotics. Many strains also express multiple drug efflux pumps and carbapenemases making *P. aeruginosa* one of the major causes of nosocomial infections in the developed and developing world. One class of molecule that readily translocate across the impervious outer membrane of *P. aeruginosa* to deliver a cytotoxin are S-type pyocins, which are 40-90 kDa protein bacteriocins made by *P. aeruginosa*. Indeed, a recent survey showed that >85% of *P. aeruginosa* strains encode nuclease-type pyocins within their genomes (Sharp *et al*, 2017) hinting at the importance of these protein antibiotics to inter-strain competition.

PyoS5 delivers a pore-forming domain across the outer membrane to depolarize the cell while PyoS5-producing cells are protected against the action of the toxin by ImS5, a small membrane localized immunity protein (Ling *et al*, 2010). Previous work has shown that PyoS5 binds the lipopolysaccharide (LPS)-anchored common polysaccharide antigen (CPA), which is identical across *P. aeruginosa* strains (McCaughey *et al*, 2016a) and is a major surface antigen in cystic fibrosis isolates (Lam *et al*, 1989), and that PyoS5 susceptibility depends on the ferric pyochelin transporter FptA (Elfarash *et al*, 2014). Here, we delineate how PyoS5, by parasitizing FptA and CPA in the outer membrane and in conjunction with proton motive force (PMF)-linked TonB1 in the inner membrane, delivers its cytotoxic domain into *P. aeruginosa*.

## Results and Discussion

### The structure of PyoS5 reveals a novel domain architecture underpins outer membrane transport in *P. aeruginosa*

S-type pyocins (which we simply refer to as pyocins) belong to a broad group of protein bacteriocins that includes colicins which kill *E. coli* as well as bacteriocins that target other Gram-negative bacteria, such as *Klebsiella pneumoniae*, *Serratia marcescens* and *Yersinia pestis*. Colicins, like pyocins, exploit the PMF to translocate through the cell envelope to deliver a cytotoxic domain, typically a pore-forming domain or a nuclease that cleaves DNA, rRNA or tRNA (Papadakos *et al*, 2012). Also like colicins, pyocins are multidomain toxins, their constituent domains associated with binding outer membrane receptors and the import process itself. There are currently several structures for intact colicins in the protein data bank (PDB) but only two for pyocins, pyocin PaeM and L1 (McCaughey *et al*, 2014; Barreteau *et al*, 2012). However, pyocins PaeM and L1 are atypical amongst the bacteriocins due to their small sizes (14 kDa for PaeM and 28 kDa for L1 compared to >50 kDa for most pyocins). Consequently, we know very little about the structural biology of typical pyocins found in *P. aeruginosa* genomes. Structural data are important to understanding bacteriocin uptake mechanisms, especially since the domain arrangement of pyocins is different to that of colicins. The receptor-binding domains are centrally located in colicins and their membrane translocation domains are at the N-terminus whereas in pyocins the order is reported to be reversed (Sano *et al*, 1993). This change in relative domain orientation would mean a fundamental difference in how these molecules transport across the outer membrane. We therefore set out to determine the crystal structure of PyoS5 and to define the functionality of its constituent domains.

PyoS5 was expressed and purified from *E. coli* cells (see Materials & Methods). The 57-kDa toxin was monomeric in solution and active against *P. aeruginosa* strains at sub-nanomolar concentrations (Figure S1). The protein crystallized in the P2_1_ space group and the structure was solved by a combination of single wavelength anomalous diffraction and molecular replacement to a resolution of 2.2 Å (Figure 1A, Supplementary Table S1 and Materials and Methods). The first 39 residues were absent from the final model, presumed unstructured which we refer to below as the disordered region. Otherwise, continuous electron density was observed for the entirety of the remaining protein sequence (residues 40-498). The structure shows that PyoS5 is an elongated, α-helical protein measuring 36 Å on the short axis and 195 Å on the long axis. Colicins are similarly long proteins and have disordered N-termini (Soelaiman *et al*, 2001; Wiener *et al*, 1997; Johnson *et al*, 2017). The extended conformation was confirmed by small angle X-ray scattering (SAXS) data; 93% of the modelled PyoS5 residues were within the SAXS envelope (Figure S2). Also similar to colicins is the prevalence of α-helical structure in PyoS5. PyoS5 contains 17 helices, the high preponderance of helical structure likely reflecting the need to forcibly unfold the toxin during transport into a cell and the lower forces known to be required for unfolding helices relative to β-sheets (reviewed in (Brockwell *et al*, 2005)).

**Figure 1:**
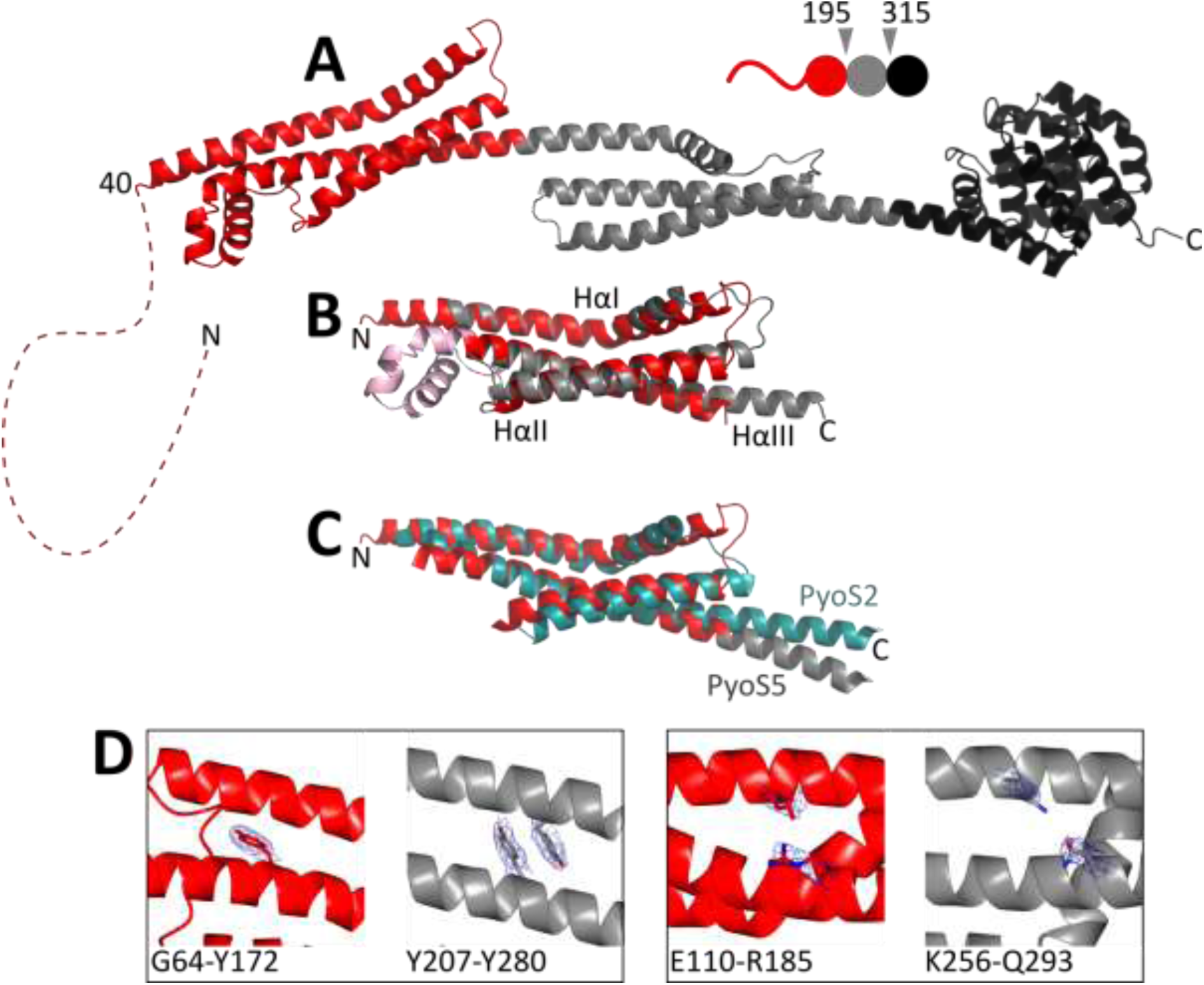
Crystal structure of PyoS5. (A) The 2.2 Å crystal structure of PyoS5 (residues 40 to 195). The first kTHB domain is in *red* (residues 196-315), the second kTHB is in grey (residues 316-505) and the pore-forming domain is in *black*. Residues 2 to 39 are not resolved and represented (to scale) by a red dashed line. (B) Structural alignment of PyoS5_40-196_ (*red*) and PyoS5_194-315_ (*grey*), RMSD 2.5 Å. Residues 123-162 (*pink*) are not conserved in PyoS5_194-315_ and were excluded from the alignment. (C) Structural alignment of PyoS5 kTHB domains (*red* and *grey*) with that from PyoS2 (*teal*), RMSD 4.1 Å. PyoS5 residues 2-213 are shown, with 123-156 excluded and PyoS2 residues 46-206 are shown, with 124-151 excluded. (D) Interactions within domain 1 (*red*) and domain 2 (*grey*) are not conserved, as illustrated by exemplary interactions shown. Electron density is shown, cut-off 1 σ.

The structure of PyoS5 is comprised of three ordered domains (Figure 1A). The C-terminal domain (domain 3; residues 315-498) has the canonical ten-helical bundle fold of a pore-forming domain found in colicins (Cascales *et al*, 2007), which is consistent with the killing activity of PyoS5 (Ling *et al*, 2010). Previous studies have highlighted that the protective immunity proteins of pore-forming domains within colicins fall into two sub-groups although the functional significance of this is unclear. Immunity proteins against colicins A, B and N – the so-called A-Type - have four transmembrane helices while those against E1, Ia and K – the so-called E1-type – have three (Cascales *et al*, 2007). Based on the predicted number of transmembrane helices of its immunity protein the pore-forming domain of PyoS5 belongs to the E1-type (Parret & De Mot, 2000). Through detailed structural comparisons of all pore-former domains with that of PyoS5 we identified a clear structural difference between the pore-forming domains of the A- and E1-groups (Figure S3). Specifically, this difference relates to the positioning of helices 1 and 5 of the domain with respect to each other; in A-type structures, helix 1 is positioned close to the centre of the domain, pushing out helix 5, while in E1-type structures, helix 5 is located closer to the centre of the domain. These pore-forming domain structures represent the ground state of the ionophore before depolarization of the inner membrane. We speculate the structural alterations evident in the A and E1-groups may reflect differences in the way each class of pore-forming domain is recognised by its particular type of immunity protein before insertion in the bacterial inner membrane.

The other structured domains of PyoS5 are also helical bundles but of a novel fold. Domain 1 comprises residues 40-194 while domain 2 comprises 195-315. The core structural motif of each domain is a kinked three-helix bundle (kTHB). The two kTHB domains are structurally similar to each other (superposition root-mean-square deviation (RMSD), 2.5 Å) but share little sequence identity (∼12%) (Figure 1B). Each kTHB domain is composed of a kinked helix I connected to a straight helix II by a loop. Helix II packs against both helix I and a third straight helix, helix III. The connection between helices II and III varies between the two copies of the fold. In domain 1, this connection is composed of three short helical turns while in domain 2 it is a loop. The other striking feature of the kTHB structural motif is that the third helix from each domain extends into the next domain of the pyocin; helix III of domain 1 extends over 90 Å into domain 2, where it forms helix I, while helix III of domain 2 extends over 90 Å to the pore-forming domain of the toxin. The kTHB fold is stabilised predominantly by hydrophobic interactions mediated by aliphatic amino acid side chains and, in one instance, aromatic stacking (Tyr207-Tyr280, domain 2) (Figure 1D). None of these stabilising interactions are conserved.

Recently, White et al reported the structure of the N-terminal domain of the nuclease pyocin PyoS2 bound to the outer membrane protein FpvAI (White *et al*, 2017). We found by structural superposition that the kTHB domain 1 of PyoS5 is structurally similar to this domain of PyoS2 (Figure 1C) and sequence similarity of 75% between the second domains of PyoS5 and PyoS2 suggest similar structures here as well (Figure S4). Sequence similarities of domains in pyocins S1, SD1, SD2, S3, SD3 and S4 to the kTHB domain also suggest these are common among pyocins (Figure S4). The structural superposition of the PyoS5 and PyoS2 kTHB domains, without the small helices connecting helix II and helix III in PyoS5, has an RMSD of 4.1 Å over 128 residues (Figure 1C).

We conclude that PyoS5 is an elongated bacteriocin comprising a disordered region at its N-terminus, two kTHB domains, which is a common structural platform for protein bacteriocins targeting *P. aeruginosa*, and a C-terminal pore-forming domain. We next set out to ascribe functions to each of the domains/regions of PyoS5 that transport the pore-forming domain into *P. aeruginosa* cells.

### Functional annotation of PyoS5 domains

We expressed and purified truncations of PyoS5 that removed one or more domains/regions. These included PyoS5_1-315_, in which the pore-forming domain was removed, PyoS5_1-196_, in which both domain 2 and the pore-forming domain were deleted, and PyoS5_194-315_, which only contained domain 2. The constructs were folded, as determined by circular dichroism spectroscopy, and their thermal melting temperatures largely recapitulated those found in intact PyoS5 (Figure S5).

We first analysed the capacity of PyoS5 and the various deletion constructs to bind CPA in isothermal titration calorimetry (ITC) experiments. Heats of binding were observed for PyoS5_1-315_ and PyoS5_194-315_ but not PyoS5_1-196_ (in 0.2 M Na-phosphate buffer pH 7.5) (Figure 2A-C, Supplementary Table S2). From these experiments, equilibrium dissociation constants (K_d_s) of 0.6 μM for PyoS5_1-315_ and 0.3 μM for PyoS5_194-315_ were obtained, similar to that reported previously for intact PyoS5 binding CPA (McCaughey *et al*, 2016a). When polysaccharides derived from *P. aeruginosa* PAO1 Δ*rmd* were used (that do not contain CPA) no binding to PyoS5_194-315_ was detected (Figure 2C). These results demonstrate that the CPA binding activity of PyoS5 resides within domain 2, and that the CPA-binding function is not a conserved feature of the kTHB fold. Pyocins S2 and SD3 have also been shown previously to bind *P. aeruginosa* CPA sugars (McCaughey *et al*, 2016a). Sequence alignments show that each has a domain equivalent to that of domain 2 of PyoS5. Indeed, the level of sequence identity across this region (39 %) is far greater than that between the two kTHB domains of PyoS5. Moreover, over half of the 45 identical residues shared between pyocins S2, SD3 and S5 form a grooved surface that runs perpendicular to the long axis of PyoS5 (Figure S4+S6). We infer that this conserved groove is the CPA binding site in these different pyocins, each of which nevertheless delivers a different cytotoxic domain into *P. aeruginosa*.

**Figure 2:**
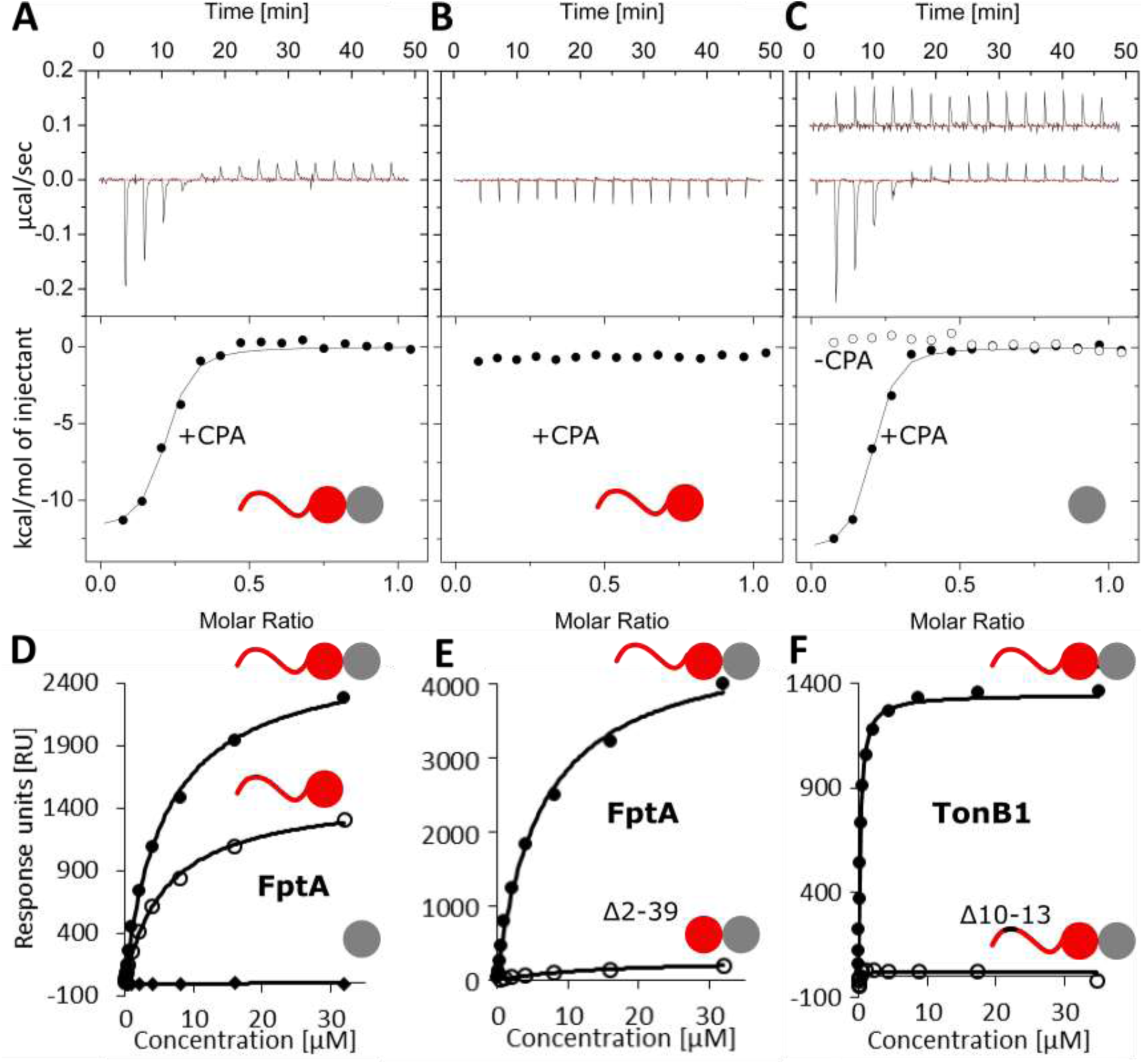
kTHB domain 2 binds CPA, kTHB domain 1 binds FptA and the N-terminal disordered region binds TonB1. (A) ITC data for PyoS5_1-315_ titrated into *P. aeruginosa* PAO1 LPS-derived polysaccharide containing CPA and OSA (closed circles) and (B) PyoS5_1-196_ titrated into *P. aeruginosa* PAO1 LPS-derived polysaccharide. (C) ITC data for PyoS5_194-315_ titrated into *P. aeruginosa* PAO1 LPS-derived polysaccharide. PyoS5_194-315_ titrated into *P. aeruginosa* Δ*rmd* LPS-derived polysaccharide containing OSA only (open circles). (A-C) K_d_s and concentrations can be found in Table 3. All ITC experiments were performed in duplicate in 0.2 M Na-phosphate buffer pH 7.5 at 25 °C, one repeat is shown. Data were corrected for heats of dilution by subtracting the average of the last five injections and fit to a model of single-site binding. (D) SPR data for FptA (0.03-32 μM) binding to PyoS5_1-315_ (*closed circles*), PyoS5_1-196_ (*open circles*) or PyoS5_194-315_ (*diamonds*). (E) SPR data for FptA (0.03-32 μM) binding to PyoS5_1-315_ (*closed circles*) or PyoS5_1-315_ Δ2-39 (*open circles*). (F) SPR data for TonB1 (0.009-35 μM) binding to PyoS5_1-315_ (*closed circles*) or PyoS5_1-315_ Δ10-13 (*open circles*). (D-F) One of three repeats is shown. All experiments were performed in parallel on the same chip in HBS-OG buffer at 25 °C. All ligands were immobilized by amine-coupling and sensorgram data was extracted and fit with a 1:1 binding model. K_d_s are presented in Table 4.

PyoS5-mediated killing of *P. aeruginosa* cells requires the ferric pyochelin transporter FptA, and the central region of the toxin (residues 151-300) has been implicated in defining this specificity (Elfarash *et al*, 2014). This region corresponds largely to domain 2 in the PyoS5 crystal structure, which, as the work above indicates, is involved in CPA binding. We therefore investigated PyoS5 binding to FptA and identified the region involved. Initially, we used native mass spectrometry (MS) to verify that PyoS5 binds FptA (Supplementary Table S3). We then determined the affinity for the complex using surface plasmon resonance (SPR) where the pyocin, and various deletion constructs, were immobilized on the chip (Figure 2D, Supplementary Table S4). These experiments determined the K_d_ for the PyoS5-FptA complex as 6.5 μM (in 25 mM HEPES buffer pH 7.5, 150 mM NaCl, 1% (w/v) n-octyl-β-D-glucoside (β-OG). Upon addition of ferric pyochelin to our SPR experiments, binding of PyoS5 to FptA reduced significantly (Figure S7B), suggesting the binding sites for the pyocin and pyochelin overlap. This result was confirmed by native state mass spectrometry experiments where PyoS5 dissociated pre-formed complexes of ferric pyochelin bound to FptA (Figure S7A). We next delineated the FptA binding site in PyoS5. Deletion of domain 2 had a marginal effect on FptA binding while domain 2 alone showed no FptA binding (Figure 2D, Supplementary Table S4). Deletion of the disordered region at the N-terminus of PyoS5 (residues 2-39) had a large effect on the amount of FptA that could bind to the chip (Figure 2E), suggesting this was affecting binding. However, closer examination indicated binding was affected only two-fold (Supplementary Table S4) and that the impact of the truncation was likely due to restricted access of FptA to its binding site on domain 1 in this construct (Figure 2E, Supplementary Table S4). By contrast, when the first 13 residues of this region were deleted (PyoS5_1-315_ Δ2-9 and PyoS5_1-315_ Δ10-13) binding to FptA remained unaffected (Supplementary Table S4). We conclude that the FptA binding site in PyoS5 is predominantly localised to kTHB domain 1 with a minor contribution from its associated disorder region at the N-terminus.

All protein bacteriocins access the PMF via either the Tol or Ton systems of Gram-negative bacteria (generally referred to as group A and B toxins in the colicin literature), which they use to drive translocation across the outer membrane (Kleanthous, 2010). It has yet to be established which of these systems is contacted by PyoS5. Typically, Tol/Ton dependence is evaluated using deletion strains. We focused initially on Ton dependence since deletion strains in *P. aeruginosa* PAO6609 are available (Tol is essential in *P. aeruginosa*). *P. aeruginosa* harbours three *tonB* genes, *tonB1*, *tonB2* and *tonB3* (Zhao & Poole, 2000; Takase *et al*, 2000; Huang *et al*, 2004). PAO6609 is a derivative of *P. aeruginosa* PAO1 and so is naturally immune to PyoS5 because it harbours the ImS5 immunity gene (Hohnadel *et al*, 1986). We therefore generated a PyoS5-ColIa chimera in which the pore-forming domain of PyoS5 was substituted for that of colicin Ia to overcome this immunity. PyoS5-ColIa was active against *P. aeruginosa* PAO6609 and strains with *tonB2* and *tonB3* deleted (Figure S8). It was not possible to test the susceptibility of a *tonB1* deletion strain because the high levels of iron needed for growth of this strain diminished PyoS5-ColIa chimera susceptibility in the parent *P. aeruginosa* PAO6609, most likely due to iron-dependent down-regulation of FptA expression (Ankenbauer & Quan, 1994). We therefore resorted to direct SPR binding assays to determine if PyoS5 bound purified TonB1 *in vitro* (see Materials and Methods for further details). We found that TonB1 binds PyoS5_1-315_ with an affinity of 230 nM in SPR experiments (Figure 2F, Supplementary Table S4). Moreover, a putative 9-residue TonB box, found in TonB1-dependent transporters and bacteriocins utilizing TonB1, is also found in the N-terminal disordered region of PyoS5 (residues 6-14). Deletion of residues 10-13 abolished binding to TonB1, confirming this region as the TonB1 binding site (Figure 2F, Supplementary Table S4).

In summary, through a combination of biophysical and structural approaches we have delineated the major binding interactions of PyoS5 with the *P. aeruginosa* cell envelope. Of the two kTHB domains, domain 2 binds CPA while domain 1 binds the ferric pyochelin transporter FptA with a minor contribution by the disordered region, which in addition binds the inner membrane protein TonB1.

### Surface accumulation and energized import of fluorescently labelled PyoS5 into *P. aeruginosa* PAO1 cells

We developed a fluorescence-based import assay for PyoS5 where transport of all its domains, barring the pore-forming domain, could be visualised and where the energetics of import could be established. We replaced the pore-forming domain of PyoS5 with a C-terminal cysteine residue and labelled this residue with AlexaFluor488 (PyoS5_1-315_-AF^488^). *P. aeruginosa* PAO1 cells were used in these experiments since cytotoxic activity was not being monitored. PyoS5_1-315_-AF^488^ readily labelled *P. aeruginosa* PAO1 cells (Figure 3A). Trypsin treatment of these labelled cells, to remove surface bound PyoS5, reduced fluorescence intensity significantly (∼eight-fold), but fluorescence was still associated with cells (Figure 3A+B). Inclusion of the protonophore carbonyl cyanide *m*-chlorophenyl hydrazone (CCCP) with the trypsin treatment completely eradicated this remaining fluorescence suggesting this protected fluorescence was internalised due to the PMF (Figure 3A+B). We next generated AF^488^-labelled constructs where either domain 2 was removed (PyoS5_1-196_-AF^488^) or where only labelled domain 2 was added to cells (PyoS5_194-315_-AF^488^). Removal of the CPA-binding domain (domain 1, PyoS5_1-196_-AF^488^) decreased surface bound fluorescence in the absence of trypsin while addition of trypsin still revealed internalised fluorescence (Figure 3C). PyoS5_194-315_-AF^488^ (domain 2 construct) on the other hand labelled cells much less efficiently (likely due to its weak binding of CPA on the surface) and all this fluorescence was trypsin sensitive, suggesting no internalisation (Figure 3C).

**Figure 3:**
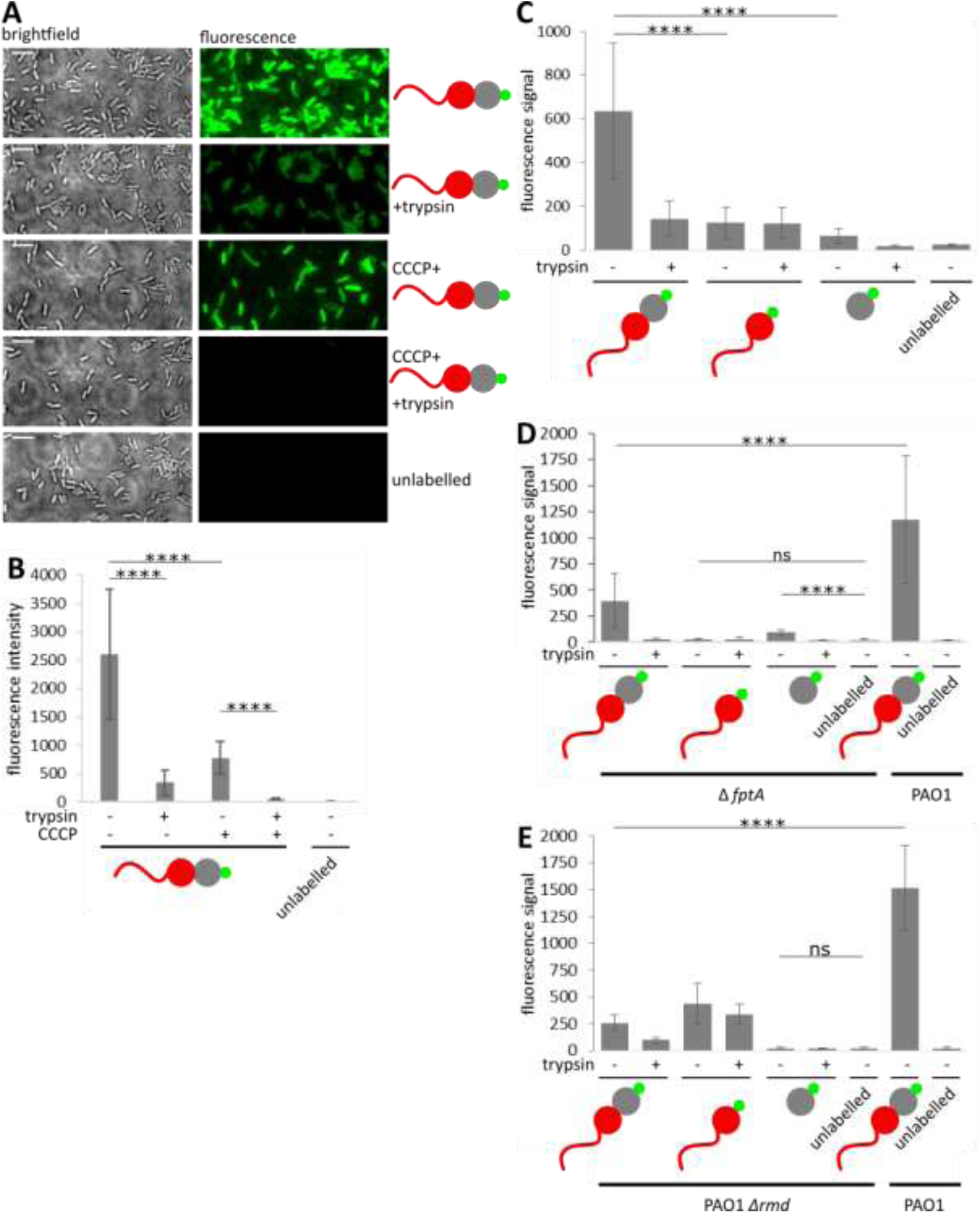
CPA accumulates PyoS5 at the cell surface while FptA and TonB1 mediate import. (A) Fluorescent labelling of live *P. aeruginosa* PAO1 cells with PyoS5_1-315_-AF^488^. Additionally, the effects of depleting the PMF with CCCP before incubation with TF-AF^488^ and of trypsin treatment to remove surface exposed PyoS5_1-315_-AF^488^ after incubation with PyoS5_1-315_-AF^488^ were examined. Scale bars 5 μm. (B) Quantification of the average cell fluorescence observed under different conditions tested in A. (C) Fluorescent labelling of live *P. aeruginosa* PAO1 using PyoS5_1-315_-AF^488^, PyoS5_1-196_-AF^488^ and PyoS5_194-315_-AF^488^ with and without trypsin treatment quantified to determine the average cell fluorescence. (D) Fluorescent labelling of live *P. aeruginosa* PW8161 (Δ*fptA*) and *P. aeruginosa* PAO1 using PyoS5_1-315_-AF^488^, PyoS5_1-196_-AF^488^ and PyoS5_194-315_-AF^488^ with and without trypsin treatment was quantified to determine the average cell fluorescence. (E) Fluorescent labelling of live *P. aeruginosa* PAO1 Δ*rmd* and PAO1 using PyoS5_1-315_-AF^488^, PyoS5_1-196_-AF^488^ and PyoS5_194-315_-AF^488^ with or without trypsin treatment was quantified to determine the average cell fluorescence. (A-E) **** indicates a p-value below 0.0001 in Student’s t-test.

Repeating these assays with *P. aeruginosa* PAO1 *ΔfptA* cells or using PyoS5_1-315_ Δ10-13-AF^488^, in which part of the TonB1 binding site (residues 10-13) was deleted, showed that trypsin-protected fluorophores (i.e. imported molecules) were no longer detected, consistent with PMF/TonB1-dependent import of PyoS5 across the outer membrane via FptA (Figure 3D and S9). Finally, import assays were conducted using *P. aeruginosa* PAO1 *Δrmd* cells, which lack CPA. Surface-associated fluorescence of PyoS5_1-315_-AF^488^ and susceptibility to PyoS5ColIa was much reduced in these cells, consistent with CPA being required for surface accumulation of PyoS5, but imported fluorescence in a domain 2 deletion was unaffected (Figure 3E and S10).

In summary, our fluorescence assays suggest that import of PyoS5 occurs in two stages. Initial binding to CPA via the central kTHB domain leads to accumulation on the surface of *P. aeruginosa*. Thereafter, the first kTHB domain of the pyocin binds FptA in the outer membrane, which also likely acts as the translocation channel, allowing contact between the disordered TonB1 binding site of PyoS5 with TonB1 in the inner membrane and PMF-driven import of the toxin (model presented below).

### Engineering pyocin susceptibility in *E. coli*

As with most bacteriocins, pyocins are specific for a subset of strains, in this case from *P. aeruginosa*, which reflects the array of cell envelope interactions required for import. Yet common principles are beginning to emerge suggesting generic import mechanisms may apply for all Gram-negative bacteria that exploit protein bacteriocins. We therefore devised a test of this hypothesis by engineering *E. coli* susceptibility towards PyoS5 utilising our current understanding of its import pathway.

Our strategy was based on first determining if the pore-forming domain of PyoS5, if imported, could kill *E. coli* cells and then engineering the minimal requirements into *E. coli* in order for PyoS5 to be recognised and transported. A similar strategy was reported by Bosak et al (2012) where *E. coli* was engineered to be susceptible to a bacteriocin specific for *Yersinia kristensenii* (Bosák *et al*, 2012). In the present work, we first showed that a chimera of the PyoS5 pore-forming domain fused to the C-terminus of the colicin B translocation and receptor-binding region (replacing colicin B’s own pore-forming domain) was cytotoxic against *E. coli* BL21 (DE3) cells. We next challenged *E. coli* BL21 (DE3) cells expressing *P. aeruginosa* FptA but saw no PyoS5 killing (Figure 4). Rationalising that *E. coli* TonB may not be recognising the TonB1 binding sites (Ton boxes) of FptA and/or PyoS5, we also expressed, in *E. coli* BL21 (DE3) cells expressing FptA along with a chimera of *E. coli* TonB (TonB_1-102_) fused to *P. aeruginosa* TonB1_201-342_. In this chimera, TonB-B1, the C-terminal domain and periplasmic regions of TonB are those from *P. aeruginosa* but the transmembrane domain that associates with TonB’s partner proteins ExbB and ExbD are those from *E. coli*. Under these conditions, *E. coli* became sensitized to PyoS5-mediated killing (Figure 4). To determine the generality of this cross-species killing, we expressed the *fpvAI* gene, which is recognised by PyoS2 and PyoS4, in *E. coli* cells expressing the *E. coli*-*P. aeruginosa* TonB-B1 hybrid. This strain was sensitive to both PyoS2 and PyoS4 but not to PyoS5 (Figure S11).

**Figure 4:**
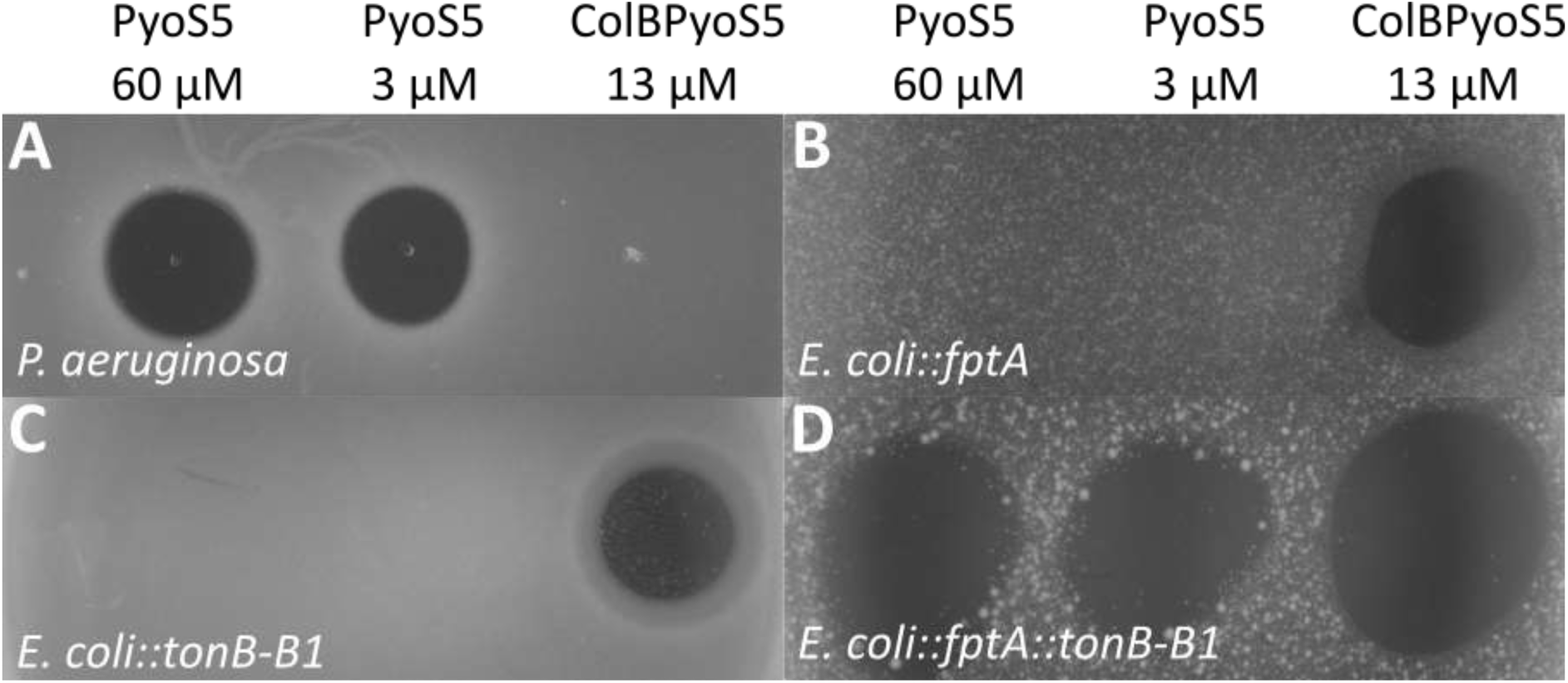
FptA and TonB1 constitute the minimal system for PyoS5-susceptibility in *E. coli*. (A) Susceptibility to PyoS5 (3 and 60 μM) was assessed for *P. aeruginosa* YHP17, (B) for *E. coli* BL21 (DE3) expressing FptA, (C) for *E. coli* BL21 (DE3) expressing TonB-B1, and (D) for *E. coli* BL21 (DE3) expressing FptA and TonB-B1. Zones of clearance are observed in all *E. coli* samples for the ColBPyoS5 (13 μM) control (B-D) and for both concentrations of PyoS5 in *E. coli* expressing FptA and TonB-B1 (D). In the *P. aeruginosa* control clearance zones were observed for PyoS5 (A).

We conclude that our engineered system is a simple means by which the import apparatus required for bacteriocins can be readily defined. Indeed, through this work we showed for the first time that PyoS4 is a TonB1-dependent bacteriocin. Importantly, our complete functional characterisation of PyoS5 demonstrates that the prevailing view of receptor-binding and translocation domains being inverted in pyocins relative to colicins is not correct. Instead, pyocins and colicins are organised in the same way, which likely explains how a pyocin can be made to work in *E. coli*. They have central receptor-binding domains (kTHB domain 2 in PyoS5) and N-terminally-located translocation domains (kTHB domain 1 and its associated disordered region). The confusion that has emerged in the field, that N-terminal domains of pyocins represent their receptor-binding domains, has arisen because pyocin interactions with their translocation channels (e.g. PyoS2 with FpvAI; (White *et al*, 2017)) can be much higher affinity than the interaction of the pyocin with its initial CPA receptor. In summary, our results suggest that the underlying mechanism by which Ton-dependent bacteriocins cross the outer membranes of the *Enterobacteriales* and *Pseudomonadales*, long thought to be unrelated, are fundamentally the same.

### Model for pyocin transport across the outer membrane of *P. aeruginosa*

White et al demonstrated recently that the N-terminal domain of PyoS2 translocates directly through FpvAI (White *et al*, 2017). The mechanism of import is analogous to that of FpvAI’s cognate siderophore ligand, ferripyoverdine; a labile portion of the transporter plug domain is removed by TonB1, allowing the TonB1 binding site (TonB box) of PyoS2 to enter the periplasm and activate import of the pyocin. Binding of PyoS2 to FpvAI is primarily through a short polyproline region that lacks regular secondary structure and mimics pyoverdine. The principal binding site of PyoS5 for FptA is domain 1 and its associated disordered region, which does not however have an equivalent polyproline sequence. Its binding to FptA is also significantly weaker than that of PyoS2 for FpvAI. For both PyoS2 and PyoS5, however, initial association with *P. aeruginosa* is by their central kTHB domains (domain 2 in PyoS5) which binds CPA embedded in the outer membrane and allows the toxin to decorate the cell surface (McCaughey *et al*, 2016a).

In Figure 5, we present a unifying model for TonB1-dependent pyocin import based on our data for PyoS5 and that presented by White et al for PyoS2 (White *et al*, 2017). CPA binding likely orients the pyocin horizontally with respect to the membrane since the predicted CPA-binding groove in PyoS5 is perpendicular to the long axis of PyoS5. This orientation assumes CPA molecules are projected vertically from the surface from their LPS anchors. After this initial surface association, we postulate that pyocins use their disordered N-terminus to find their transporter, the binding of which causes the pyocin to reorient, allowing the N-terminal kTHB domain to engage the transporter (as found in the PyoS2-FpvAI complex). Similar ‘fishing pole’ models have been proposed for receptor-bound colicins finding translocator proteins, but in these instances the receptor is generally an outer membrane protein (Zakharov *et al*, 2004). Following opening of the transporter channel by TonB1, the pyocin’s own TonB1 binding site enters the periplasm. A second PMF-dependent step then occurs in which TonB1 in conjunction with the PMF unfolds the kTHB domain of the pyocin and pulls it through the transporter. Whether this energized interaction is responsible for the entire pyocin entering the periplasm (as shown in Figure 5) or whether domain refolding in the periplasm contributes to the entry process remains to be established.

**Figure 5:**
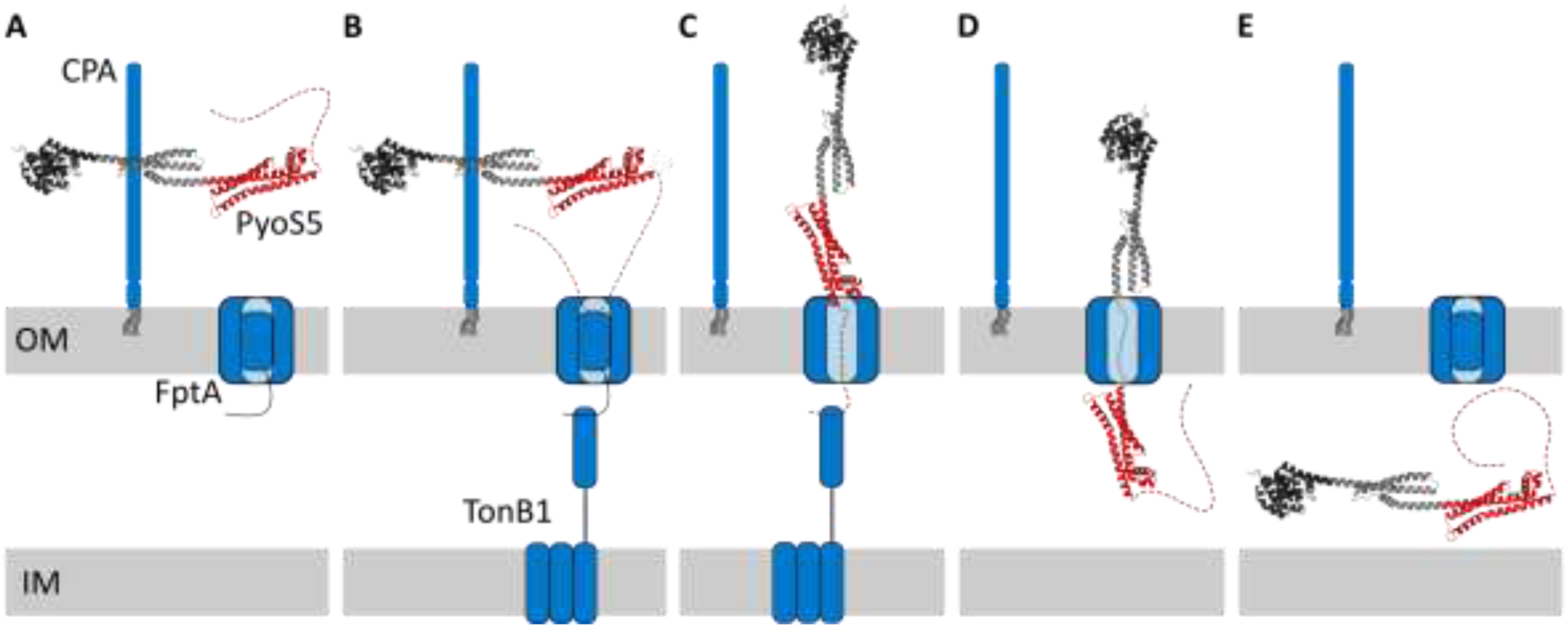
Model of PyoS5 import. (A) PyoS5 accumulates on the cell surface by binding to CPA through kTHB domain 2. (B) PyoS5 then contacts its outer membrane translocator, FptA, initially with its disordered N-terminus, and then through binding of kTHB domain 1. (C) Interactions between FptA and TonB1 possibly act to induce movement of the receptor plug domain, allowing for the unstructured N-terminus of PyoS5 to thread through the receptor and access the periplasm. Following entry to the periplasm the N-terminus of PyoS5 binds toTonB1 though the TonB-box motif. The formation of the PyoS5-TonB1 complex enables coupling to inner membrane protein targets of TonB1. (D) This coupling provides energy transduction from the PMF that facilitates translocation of PyoS5 through the outer membrane. (E) Finally, this results in PyoS5 translocation into the periplasm.

## Materials and methods

Pyochelin was synthesized as described previously (Zamri & Abdallah, 2000). Chromatography columns were purchased from GE Healthcare.

### Strains and plasmids

All bacteria (Table 2-1) were cultured in LB (10 g/L tryptone, 10 g/L NaCl, 5 g/L yeast extract, pH 7.2) at 37 °C at 120 rpm shaking unless otherwise stated. Liquid cultures were inoculated from single colonies on LB agar (1.5% (w/v)) plates. M9 medium (8.6 mM NaCl, 18.7 mM NH_4_Cl, 42.3 mM Na_2_HPO_4_, 22.0 mM KH_2_PO_4_) was supplemented with 0.4% (w/v) glucose, 2 mM MgSO_4_, and 0.1 mM CaCl_2_.

**Table 1:** Bacterial strains used in this study.

| Name | Relevant characteristics | Source |
| --- | --- | --- |
| ***E. coli*** |  |  |
| NEB5α | *fhuA2 Δ(argF-lacZ)U169 phoA glnV44 Φ80 Δ(lacZ)M15 gyrA96 recA1 relA1 endA1 thi-1 hsdR17* | New England Biolabs |
| BL21 (DE3) | *fhuA2 [lon] ompT gal (λ DE3) [dcm] ΔhsdS λ DE3 = λ sBamHI ΔEcoRI-B int::(lacI::PlacUV5::T7 gene1) i21 Δnin5* | New England Biolabs |
| TNE012 | *ompA<sup>-</sup>, ompB<sup>-</sup>, tsx<sup>-</sup>* | Professor William Cramer (Taylor *et al*, 1998) |
| ***P. aeruginosa*** |  |  |
| PAO1 | Clinical isolate | Manoil lab Washington mutant library |
| YHP17 | Clinical isolate | Professor Daniel Walker |
| PAO6609 | *met-9011 amiE200 strA pvd-9* | Professor Iain Lamont (Hohnadel *et al*, 1986) |
| K1407 | *PAO6609 tonB1<sup>-</sup>* | Professor Iain Lamont (Poole *et al*, 1996; Zhao & Poole, 2000) |
| K1408 | *PAO6609 tonB2<sup>-</sup>* | Professor Iain Lamont (Zhao & Poole, 2000) |
| MS231 | *PAO6609 tonB3<sup>-</sup>* | Professor Iain Lamont (Shirley & Lamont, 2009) |
| MS233 | *PAO6609 tonB2<sup>-</sup>, tonB3<sup>-</sup>* | Professor Iain Lamont (Shirley & Lamont, 2009) |
| PW8161 | *PAO1 fptA<sup>-</sup>* | Manoil lab Washington mutant library |
| PAO1 *Δrmd* | *CPA deficient* | Professor Cezar Khursigara (Rocchetta *et al*, 1998) |

**Table 2:**
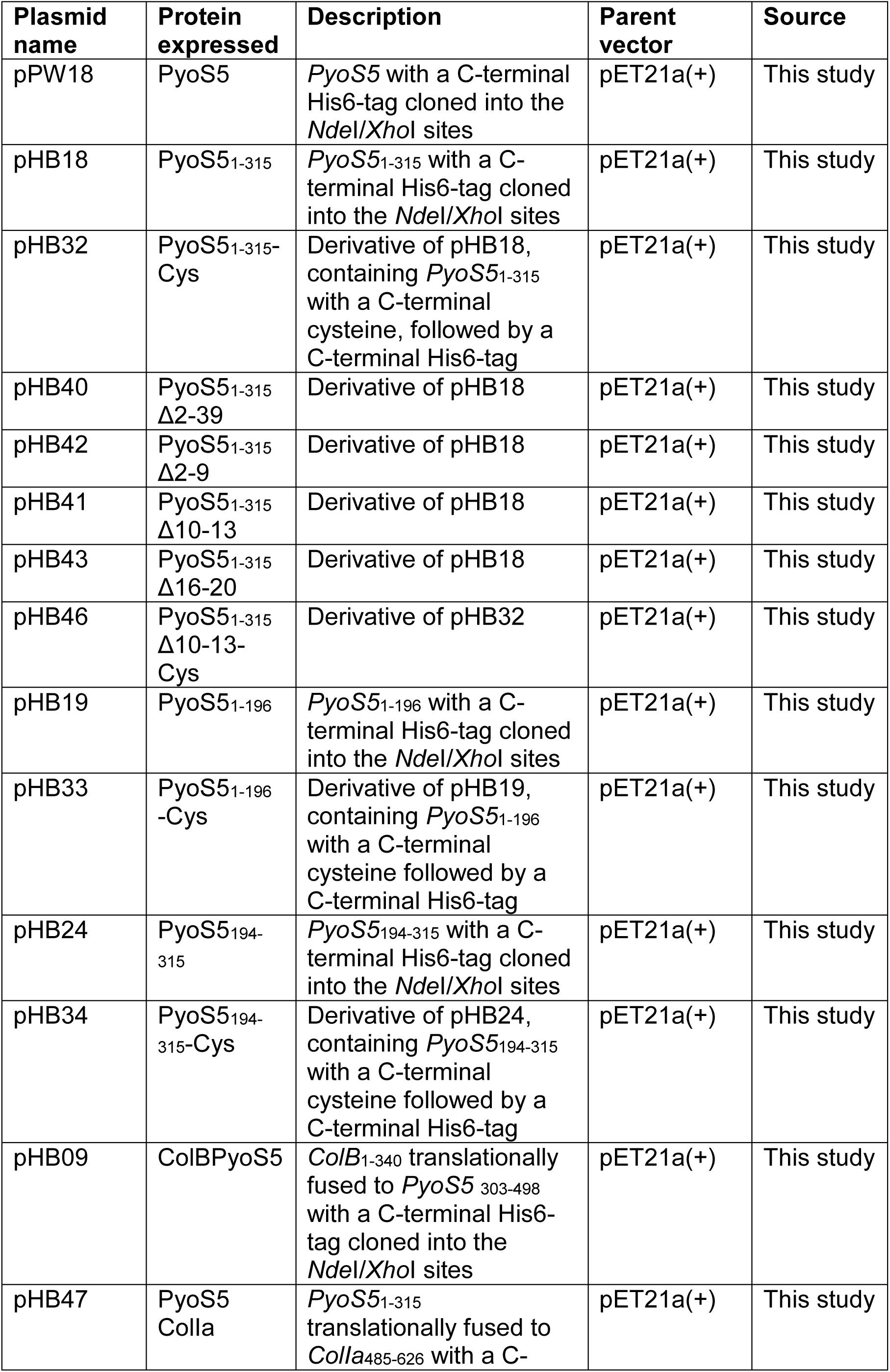
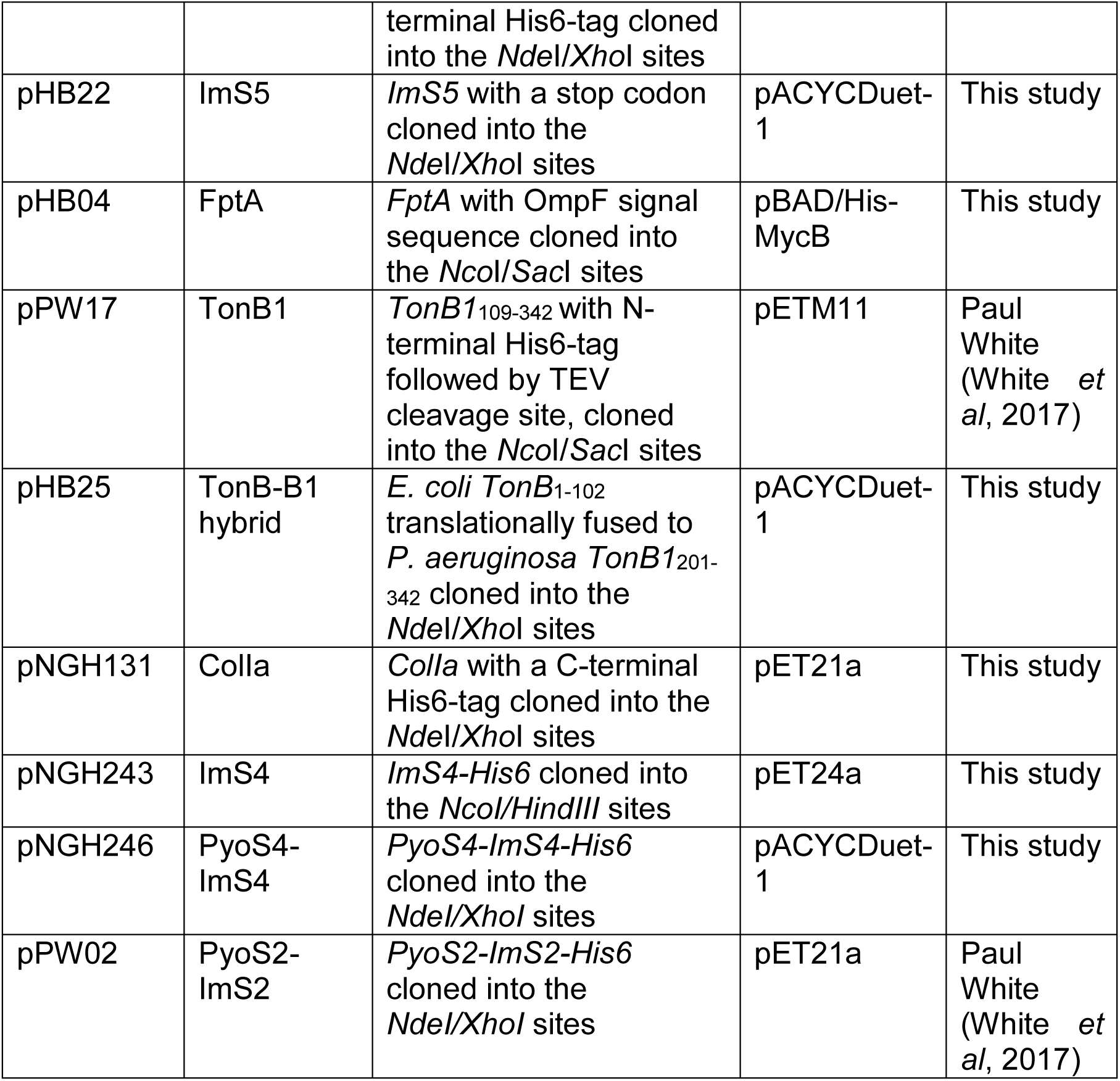
Expression plasmids used in this study.

### Molecular biology

Genes were amplified from genomic DNA or synthesized by Genewiz. Plasmids were created by restriction enzyme digest and ligation or quick-change mutagenesis. Chemically competent *E. coli* NEB5α and BL21 (DE3) were purchased from NEB. Antibiotics were used at the following final concentrations; ampicillin, 100 μg/mL, kanamycin and gentamicin, 50 μg/mL, all from stock solutions in water; chloramphenicol at 37 μg/mL and tetracyclin at 10 μg/mL from stock solutions in ethanol.

### Expression and purification of bacteriocins

PyoS5 and its derivatives, as well as ColBPyoS5, PyoS5ColIa, PyoS2, PyoS4 and ColIa were expressed heterologously from *E. coli* BL21 (DE3) for 3 h at 37 °C or overnight at 20°C while shaking at 120 rpm. For constructs containing the PyoS5 pore-forming domain (amino acid residues 315-498) the cells were co-transformed with pHB22 which carries the ImS5 immunity protein, for increased yield. The bacteria were harvested at 5050 g for 15 min at 10 °C, resuspended in binding buffer (0.5 M NaCl, 20 mM Tris-HCl pH 7.5) and sonicated on ice. They were then centrifuged at 12500 g for 20 min at 4 °C, filtered through a 0.45 μm syringe filter, loaded onto a 5 mL HisTrap-HP-column equilibrated in binding buffer and eluted by gradient elution using elution buffer (binding buffer + 0.75 M imidazole). The protein was then dialyzed into size exclusion buffer (150 mM NaCl, 20 mM Tris-HCl pH 7.5) using a 12-14 kDa molecular weight cut off membrane (Spectra/Por, Spectrum), filtered through a 0.45 μm syringe filter and applied to a 26/60 Superdex 200 size exclusion chromatography column.

PyoS4 was expressed at 28 °C in the presence of an additional copy of ImS4 (pNGH243), and purified on an S200 16/60 size exclusion column.

Mass spectrometry indicated that all bacteriocins purified without their N-terminal methionines, with exception of PyoS5_1-315_ Δ2-20, PyoS5_194-315_ and PyoS5_194-315_-Cys.

### Expression and purification of TonB1 soluble fragments

The TonB1 construct was purified by HisTrap-HP-column as described for PyoS5 and then incubated in 300 mM NaCl, 50 mM Tris-HCl pH 7.0 with 0.07 mg/mL 6-His-TEV (tobacco etch virus-protease) at RT for 4.5 h. TonB1 was then purified by affinity chromatrography on a HisTrap-HP column and by size exclusion chromatography on a 26/60 Superdex 200 column.

### Expression and purification of FptA

FptA purification was modelled after a previous BtuB purification protocol (Housden *et al*, 2005). FptA was expressed heterologously from *E. coli* TNE012 at 37 °C while shaking at 120 rpm in LB, and upon reaching OD_600_ 0.6 induced with 0.15% (w/v) arabinose and supplemented with 0.15% w/v glucose. The bacteria were harvested as described for PyoS5 and resuspended in 10 mM Tris-HCl pH 8.0, 0.25% (w/v) lithium diiodosalicylic acid (LIS), sonicated as described for PyoS5 and centrifuged at 4000 g for 20 min at 4 °C. The supernatant was collected, the pellet resuspended in fresh buffer and centrifuged again. Both supernatants were ultra-centrifuged at 200 000 g for 45 min at 4 °C. The pellet was homogenized in 10 mM Tris-HCl pH 8.0, 0.25% (w/v) LIS, 2% (v/v) Triton X-100 and ultra-centrifuged again. The resulting pellet was homogenized in 10 mM Tris-HCl pH 8.0 and ultra-centrifuged again. The resulting pellet was homogenized in 10 mM Tris-HCl pH 8.0 + 2% (w/v) β-OG, 5 mM ethylenediaminetetraacetic acid (EDTA) and ultra-centrifuged again. FptA was purified from the supernatant by anion exchange chromatography. A 5 mL HiTrap DEAE FF column was equilibrated in buffer E (50 mM Tris-HCl pH 7.5, 1% (w/v) β-OG, 5 mM EDTA) and gradient eluted with buffer F (buffer E + 1 M LiCl). This was followed by 16/60 Sephacryl 300 size exclusion chromatography in buffer E and anion exchange chromatography on a Mono Q 4.6/100 PE column in buffer E, with gradient elution with buffer F.

### Protein quantification

All protein concentrations were measured using absorbance at 280 nm which was converted to concentration using the sequence based predicted molar extinction coefficient (ExPASy ProtParam). The presence of scattering impurities, such as protein aggregates, was checked for by measuring the absorbance at 320 nm. All protein masses were confirmed by denaturing electrospray ionoization (ESI) mass spectrometry (MS), performed on proteins diluted in formic acid.

### Pyocin cytotoxicity assays

*P. aeruginosa* YHP17 were grown to OD_600_ 0.6 and 200 μL of the culture were mixed with melted, 50 °C warm, soft LB agar (0.75% (w/v) agar) and poured over an LB agar plate. Once the plate had set, 2.5 μL of each bacteriocin concentration were spotted onto the plate. The plates were left to dry and then incubated at 37 °C overnight.

### LPS-derived polysaccharide isolation

LPS-derived polysaccharides were isolated as described previously (McCaughey *et al*, 2016a). Briefly, 1 L of cells were grown for 20 h at 37 °C, pelleted at 6000 g for 20 min and resuspended in 10 mL 50 mM Tris pH 7.5, 2 mg/mL lyosozyme, 0.5 mg/mL DNAse I. Cells were lysed by sonication, as described for PyoS5 isolation, the lysate incubated for 30 min at RT and then 0.2 mM EDTA added. An equal volume of aqueous phenol was then added and the mixture heated for 20 min at 70 °C with mixing. The solution was incubated on ice for 30 min, centrifuged at 7000 g for 20 min and the aqueous upper layer was extracted.

0.05 mg/mL proteinase K was added and the solution dialysed overnight against 5 L dH_2_O, followed by dialysis against 5 L fresh dH_2_O for 5 h. LPS was pelleted by ultracentrifugation for 1 h at 100 000 g and the pellet resuspended in 10 mL dH_2_O. The suspension was heated at 60 °C for 30 min, then acetic acid added and the mixture heated at 96 C for 1.5 h. Lipid A was pelleted by centrifugation at 13 500 g for 3 min and the supernatant, which contains the polysaccharide was extracted with 10 mL chloroform. The aqueous phase was then lyophilized.

### Biophysical methods

**Native mass spectrometry** was performed in 100 mM ammonium acetate buffer with exception of TonB1 which was analysed in 200 mM ammonium acetate buffer.

**SPR** was performed on a Biacore T200 instrument. A Series S Sensor Chip CM5 (GE LifeScience) was docked and primed into HBS-OG buffer (25 mM HEPES pH 7.5, 150 mM NaCl, 1% (w/v) β-OG). This buffer was used as a running buffer for all SPR experiments.

For amine coupling using the Amine Coupling kit (GE Healthcare), ligand proteins were desalted into immobilization buffer (25 mM potassium phosphate pH 7.5, 50 mM NaCl) and diluted 10-fold in 10 mM sodium acetate pH 5.0 (GE LifeScience).

For thiol coupling using the Thiol Coupling kit (GE Healthcare), ligand proteins were incubated with 10 mM (dithiothreitol) DTT for 2 h, then desalted into immobilization buffer diluted 10-fold in 10 mM sodium acetate pH 5.0 (GE LifeScience) immediately before immobilization.

Analyte proteins were desalted into HBS-OG buffer before application. The contact time for SPR was set to 120 s, the dissociation time to 600 s and the flow rate to 30 µL/min. Lower analyte concentrations were applied first.

**ITC** was performed using a MicroCal iTC200 instrument at 25 °C in 0.2 M sodium phosphate buffer pH 7.5. Proteins in the syringe were at 150 µM, polysaccharides in the cell at 7 mg/mL which estimated to be 30 µM based on a molecular weight of 10 kDa and the assumption that CPA constitutes 5% of the LPS polysaccharides. The data were fitted to a one binding site model in Microcal LLC Origin software. As the CPA concentration is estimated, the observed stoichiometry is unlikely to be correct, while ΔH, ΔS and K_d_ are unaffected by the analyte concentration. Errors reported in the text are standard deviations of the average of two experiments.

**SAXS** data were collected at the B21 Beamline at Diamond Light Source proteins following in-line size exclusion chromatography on a Superdex 200 column and processed using Scatter and Atsas (Rambo, 2019; Petoukhov *et al*, 2012). Guinier approximation analysis and P(r) distributions were determined using Scatter. Dummy atoms were fit using multiple parallel runs of DAMMIF (Franke & Svergun, 2009) and refined using DAMMIN (reference). Bead models were converted to maps using Situs (Wriggers, 2012) and structures fit into the envelopes using Chimera (Pettersen *et al*, 2004). CRYSOL from the Atsas suite was used to generate the theoretical curve of the crystal structure and to fit it to the SAXS data.

**Circular dichroism.** Proteins were analysed at 0.1 mg/mL in 10 mM potassium phosphate buffer pH 7.5, 20 mM NaCl using a Jasco J-815 Spectropolarimeter. Spectra were measured between 260 nm and 190 nm at a digital integration time of 1 s and a 1 nm band width. Each sample spectrum was measured in quadruplicate and averaged. Molar ellipticity was calculated by subtracting the baseline from sample spectra and dividing by the molecular weight, molar concentration and pathlength in mm. Thermal melting curves for proteins were measured at 222 nm between 20 °C and 86 °C and 4-parameter sigmoidal melting curves were fit to the equation f = y_0_+a/(1+*e*^((x-x0)/b^) using non-linear regressions in SigmaPlot to determine the melting temperature (T_m_).

**Size exclusion multi angle light scattering (SEC-MALS).** Proteins were separated in 50 mM Tris pH 7.5, 150 mM NaCl using a Superdex 200 10/300 GL column and detected by a Wyatt Dawn HELEOS-II 8-angle light scattering detector and a Wyatt Optilab rEX refractive index monitor linked to a Shimadzu HPLC system.

### X-ray crystallography

Pyocin S5 was concentrated to 16 mg/mL in 25 mL Tris-HCl pH 7.5, 150 mM NaCl using a VivaSpin 20 column with a 30 kDa molecular weight cut off (Sartorius). The crystallisation screens Index (Hampton Research) and PACT, JCSG+ and Morpheus (Molecular Dimensions) were used to screen for crystals. Crystals were grown in a vapour diffusion sitting drop set up in JCSG+ screen (Molecular Dimensions) condition C7 (10% (w/v) PEG 3000, 0.1 M sodium acetate, 0.1 M zinc acetate, pH 4.5) at 18 °C. Drops contained 100 nL protein and 100 nL buffer. The cryoprotectant solution was 25 % glycerol, 10% w/v PEG 3000, 0.1 M sodium acetate, 0.1 M zinc acetate, pH 4.5 for cooling the crystals in liquid nitrogen. Diffraction data were collected at beamline ID30A-3 at ESRF at a wavelength of 0.9679 Å using an EIGER detector. We collected 225 degrees of data with 0.15-degree oscillation. Transmission was 20% and exposure time was 0.010 s.

The raw data were analysed in Dials, revealing a P2_1_ space group and yielding a 98.8% complete set of indexed diffraction spots but no anomalous signal. Molecular replacement was carried out using ColIa residues 450-624 in Phaser and yielded electron density for the pore-forming domain of PyoS5. Lack of density for the remainder of the protein indicated that the phases, obtained from ColIa, were not sufficient to build a model for the whole protein.

Improved phases were obtained from anisotropy correction of the same data set using Staraniso in AutoProc (Vonrhein *et al*, 2011; Tickle *et al*, 2019), which allowed a weak anomalous signal to be detected. The partial model from molecular replacement from Dials and the anomalous data from AutoProc were combined for MR-SAD phasing using Phaser (McCoy *et al*, 2007). An anomalous substructure containing eight metal ions was identified. Based on the type of metal present in the crystallization condition, these were assumed to be Zn^+2^. The result was additional, visible helical density beyond the pore-forming domain.

Iterations of model building into the visible helical density in Coot and refinement against the complete Dials data set in Buster version 2.10.3, resulted in a model of PyoS5. The model was optimized in Coot (Emsley & Cowtan, 2004), followed by one crystallographic refinement in Buster, followed by model optimization in Coot and one refinement in Phenix 1.12 (Adams *et al*, 2010). Up to then the whole model was treated as one TLS group. At this point four new TLS groups were created based on similar B-factors as determined in Phenix, comprising residues 40 to 212, 213 to 338, 339 to 395 and 395 to 505, respectively. This increased the R_work_ and R_free_ upon refinement indicating that the use of multiple TLS groups made the model worse. The refinement process was therefore continued with the whole model treated as one TLS group.

At the end of the model optimization and refinement the R_work_ was 0.212 and the R_free_ 0.272. MolProbitiy, (Chen *et al*, 2010), was used to validate the structure and assess its quality, resulting in a Molprobity score of 1.57. At the end of this validation process the R_work_ was 0.225 and the R_free_ 0.275. Figures of the crystal structure were created using CCP4MG (Winn *et al*, 2011) and PyMOL (Delano, 2002).

### Fluorescence microscopy

#### Fluorescent labelling of proteins

Bacteriocins were fluorescently labelled using maleimide AF488 labels via an engineered C-terminal cysteine. To reduce the cysteine, the protein was mixed in a 1 to 9 ratio with DTT to yield a concentration of 10 mM DTT and incubated for 2 h at RT. To remove aggregates, the protein was centrifuged at 16000 g for 1.5 min and the supernatant transferred to a new tube. The supernatant was then applied to a 5 mL HiTrap desalting column and desalted into 25 mM Tris-HCl pH 7.5, 100 mM NaCl, 1% (w/v) β-OG. The protein concentration was measured and immediately maleimide AF488 was added in 3-fold excess. The reaction was allowed to proceed for 1 h while mixing by rotary inversion in the dark at RT. Then the reaction was quenched by adding DTT to a final concentration of 5 mM. The solution was centrifuged and desalted as before. The absorbance was measured at 280 nm and 494 nm using a V-550 UV-Visible Spectrophotometer (Jasco). Labelling efficiency was determined as described in the manufacturers protocol (Molecular Probes Inc, 2006). All fluorescently labelled proteins used for microscopy were labelled with more than 95% efficiency.

#### Fluorescent labelling of bacteria

Coverslips were cleaned by water bath sonication at 50 °C for 15 min in 2% Neutracon (Decon) solution, washed in ddH_2_O and air dried.

Bacteria were grown over night in LB medium. 1 mL of this overnight culture were pelleted and resuspended in 10 mL supplemented M9 medium and grown until an OD_600_ of 0.6. 600 µL of this culture were used per condition. All pelleting steps were performed at 7000 g for 3 min at RT.

For CCCP treatment, CCCP was added to a final concentration of 100 µM from a 10 mM stock in DMSO to the bacteria before addition of the fluorescently labelled protein. The bacteria were incubated with CCCP while mixing by rotary inversion at RT for 5 min, while all other samples were incubated without CCCP for the same time. Fluorescently labelled protein was then added to a concentration of 1 µM and the sample incubated in the dark while mixing by rotary inversion for 20 min at RT.

For trypsin treatment, trypsin was added to a final concentration of 0.1 mg/mL immediately after the incubation with the fluorophore-labelled pyocin. The bacteria were incubated with or without trypsin at 30 °C for 1 hour at 120 rpm.

Subsequently, bacteria were washed three times in supplemented M9, where each wash consisted of pelleting the bacteria, removing the supernatant, resuspending the pellet in 50 µL by repeated pipetting (10 times) with a P20 pipette, transferring the 50 µL to a new tube with 450 µL supplemented M9, and vortexing. The bacteria were resuspended in a final volume of 30 µL. 3 µL were applied to an agar pad for microscopic analysis. Agar pads were prepared using Geneframes (Thermo Scientific) as follows. 1% (w/v) supplemented M9 agar was prepared and 190 µL pipetted into the Geneframe. Using a coverslip, the surface was flattened and excess agar removed. Once the agar was solidified the cover slip was removed, the bacterial suspension added and a new coverslip attached to the adhesive side of the Geneframe.

### Image collection

All images were collected on an Oxford Nanoimager S microscope at 100 ms exposure. For every image, 200 frames were collected and averaged. Green fluorescence (excitation: 473 nm, emission 425/50 nm) was measured at 35% laser power.

### Data analysis

In ImageJ the 200 collected frames per image were merged using the command “Z project”. Bacterial cells and background were identified in trans-illumination images using “Trainable Weka Classifier”. Regions of interest were transferred to green fluorescence images and the mean fluorescence of cells, “signal”, and background “noise” quantified. Each image contained a minimum of 15 bacterial cells. For each repeat a minimum of six images were collected per sample and three independent experiments were performed for each experiment. As a result, a minimum of 270 bacterial cells were quantified for each sample. Students t-tests were performed to determine p-values.

### Sequence and structure comparisons

Sequences were compared using NCBI BLASTn and BLASTp (Altschul *et al*, 1990), MUSCLE (Madeira *et al*, 2019), and jackhmmer (Potter *et al*, 2018). Similar structures were searched for using NCBI VAST (Madej *et al*, 2014) and eFOLD (Krissinel & Henrick, 2004).

### Data availability

The data supporting the findings of the study are available in the article and its Supporting Information or from the corresponding author upon request. The crystallography data from this publication have been deposited to the PDB database https://www.rcsb.org/ and assigned the identifier 6THK.

## Acknowledgements

We are indebted to Dr David Staunton (Molecular Biophysics Suite, Oxford) for help and assistance with biophysical measurements. We thank Professor William Cramer for providing TNE012 cells, Professor Iain Lamont for providing PAO6609, K1407, K1408, MS231 and MS233 cells, and Professor Cezar Khursigara for providing PAO1 Δ*rmd* cells. This work was supported by the Wellcome Trust through the Infection, Immunology & Translational Medicine DPhil studentship to HMB and through a Collaborative Award to CK and DW. TMW was supported by the Erasmus+ scheme of the European Commission. CVR is funded by a Wellcome Trust Investigator Award (104633/Z/14/Z), an ERC Advanced Grant ENABLE (641317) and an MRC Programme Grant (MR/N020413/1). JG acknowledges support of a Junior Research Fellowship from The Queen’s College, Oxford. *P. aeruginosa* Mutant Library strain PW8161 was created with support of grant NIH P30 DK089507.

## Author contributions

HMB, NGH and CK designed the research. HMB, EDL, JG, NGH, RK, TMW and CMAT performed the research. HMB, EDL, JG, NGH and TMW analysed data. GLAM and IJS provided essential material. HMB and CK wrote the article. HMB, EDL, NGH, DW and CK revised the article. IJS, DW, CVR and CK obtained funding.

---

**Supplementary Figure S1:**
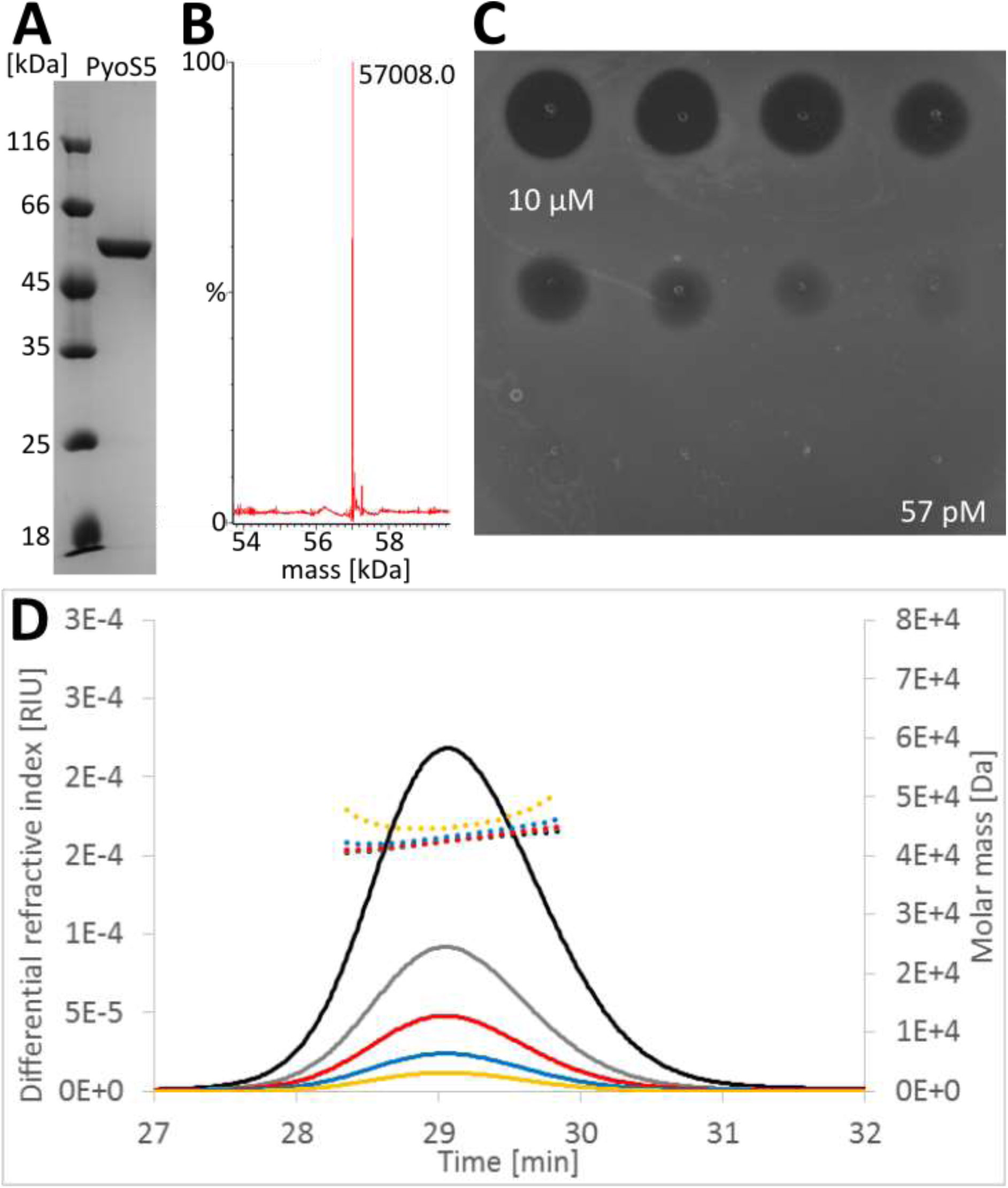
Characterisation of purified PyoS5. (A) Coomassie-stained SDS-PAGE showing purified PyoS5, compared to molecular weight standards (Fermentas unstained protein marker). (B) ESI-MS of full length PyoS5 gives a mass of 57008.00 Da, compared with a sequence-based expected mass of 58007.98 Da. (C) PyoS5 spotted on *P. aeruginosa* YHP17 in three-fold dilutions starting at 10 µM. Zones of clearance indicate growth inhibition by PyoS5. (D) SEC-MALS of PyoS5 at 9.5 mg/mL (black), 4.8 mg/mL (grey), 2.4 mg/mL (red), 1.1 mg/mL (blue) and 0.6 mg/mL (yellow) shows PyoS5 is monomeric at all tested concentrations.

**Supplementary Figure S2:**
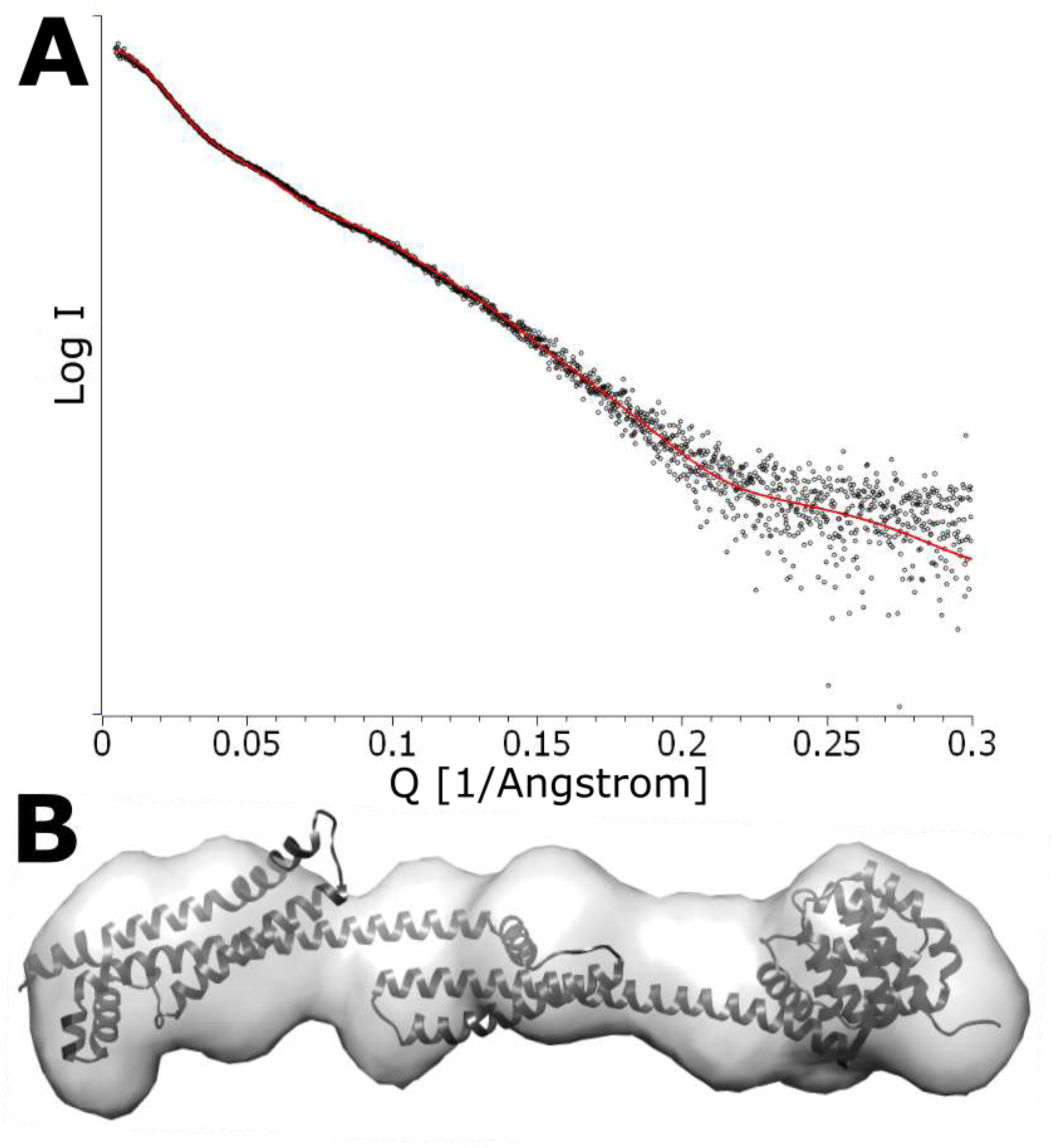
SAXS data of PyoS5. (A) SAXS of PyoS5 (black) at 5.4 mg/mL shows PyoS5 is monodispersed. The theoretical scattering curve for the crystal structure (red line) fits the experimental SAXS data well with a χ^2^ of 1.74. (B) Fit of the PyoS5 crystal structure into the PyoS5 SAXS envelope by Chimera shows 93% of atoms within the SAXS envelope.

**Supplementary Figure S3:**
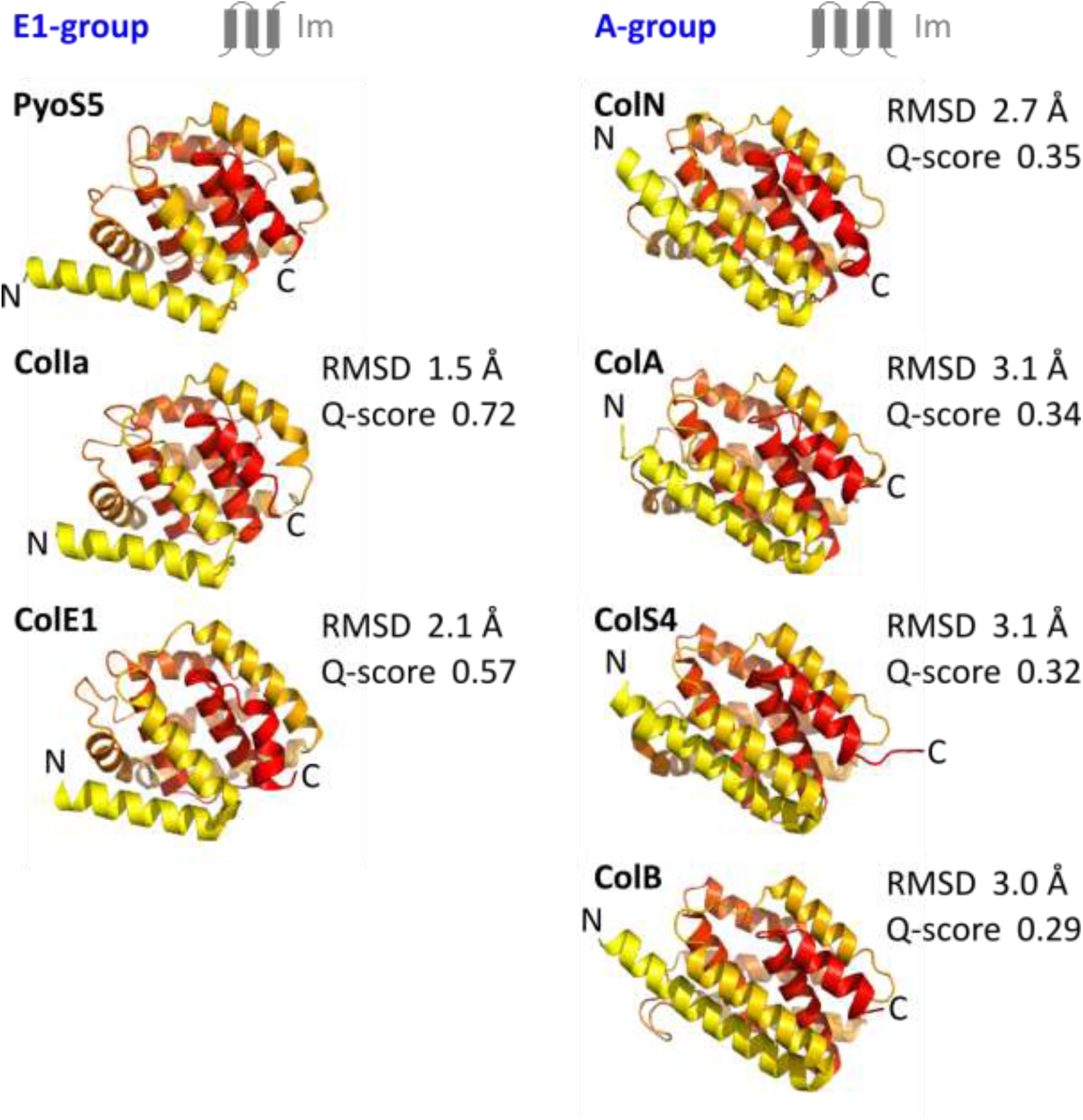
Comparison of pore forming domains from bacteriocins. All available structures of pore-forming domains were aligned and coloured in a gradient from yellow (N-terminus) to red (C-terminus). The pores are shown in order of decreasing alignment quality (Q) score. All pores in the left column have immunity proteins that pass the membrane three times, those in the right column have immunity proteins that pass the membrane four times. RMSDs, Q-scores, and immunity protein topologies are shown. The following residues were aligned to PyoS5 315-498: ColIa 448-624 (PDB 1CII), ColE1 345-522 (PDB 2I88), ColN 188-486 (PDB 1A87), ColA 393-591 (PDB 1COL), ColS4 299-499 (PDB 3FEW) and ColB 312-511 (PDB 1RH1).

**Supplementary Figure S4:**
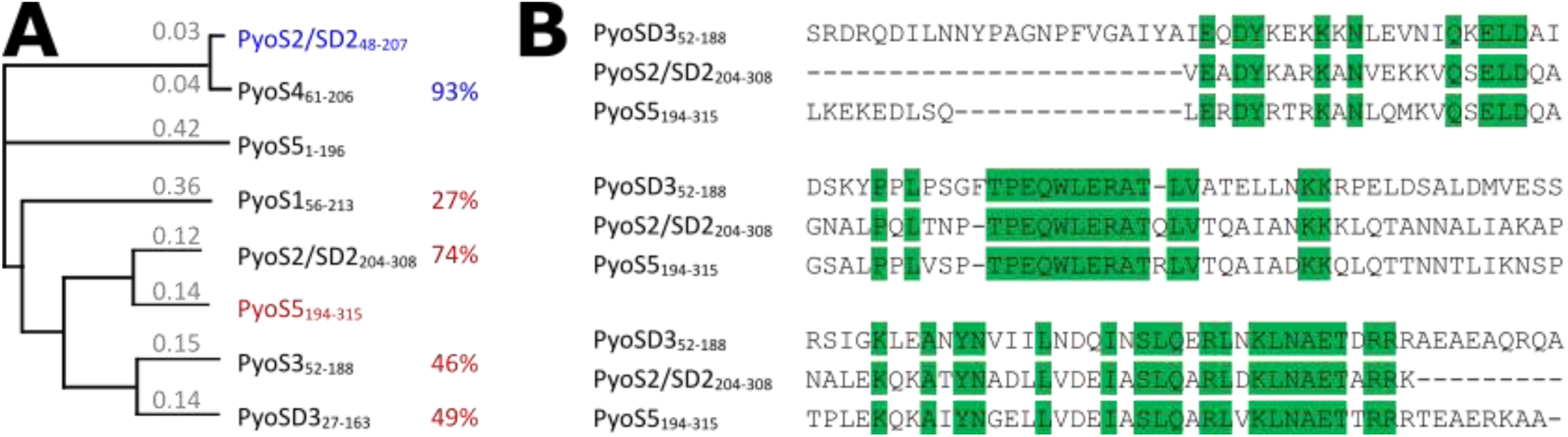
Potential kTHB domains in other pyocins. (A) Phylogentic tree of kTHB domains with branch lengths in substitutions per site (*grey*) and percent protein sequence identity to the most similar crystalized kTHB (*blue* and *red*). (B) Sequence alignment of the three kTHBs that have been shown to bind CPA. Identical residues are shown in green.

**Supplementary Figure S5:**
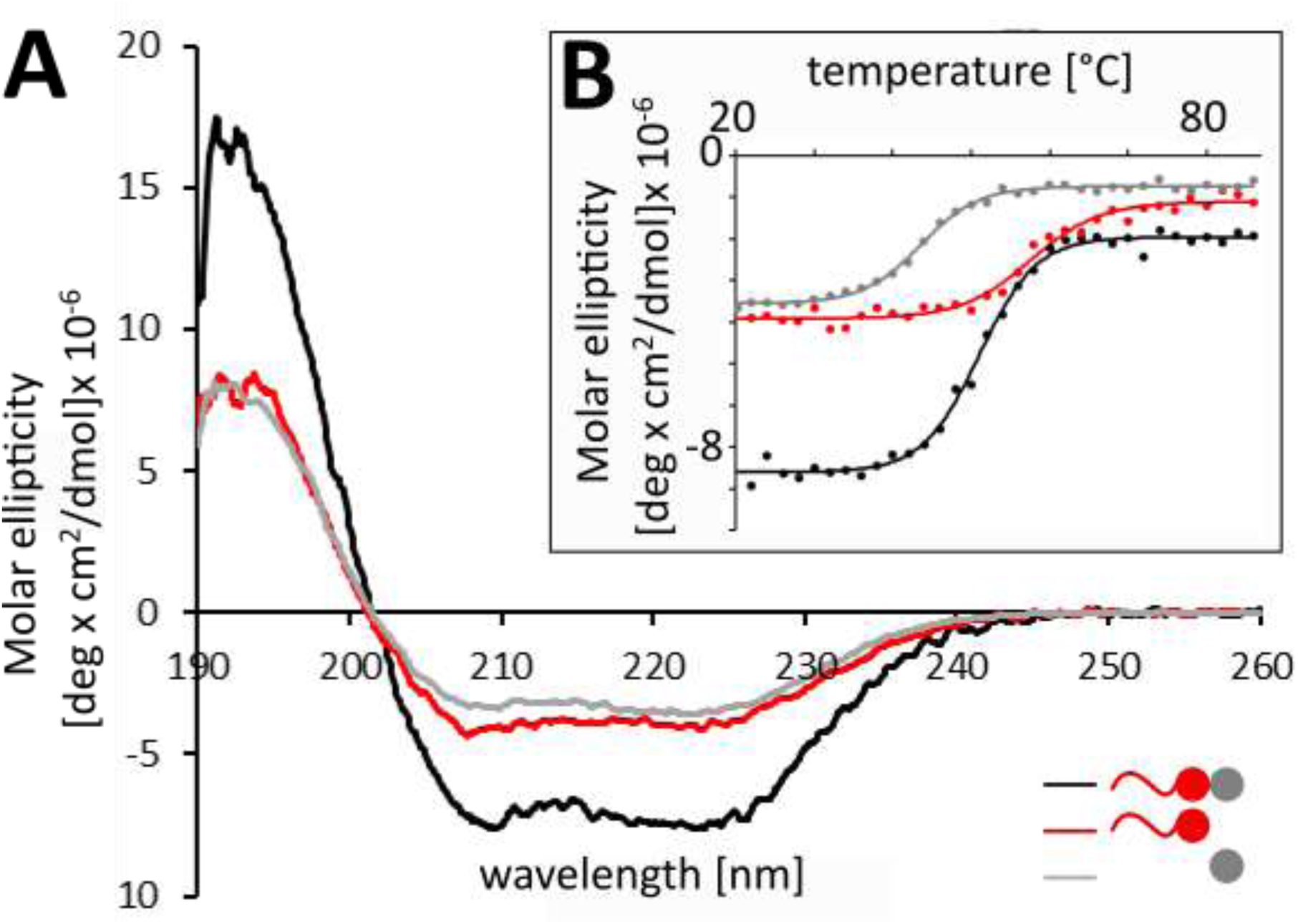
Circular dichroism (CD) spectra of PyoS5 constructs. (A) CD spectra of 0.1 mg/mL PyoS5_1-315_ (*black*), PyoS5_1-196_ (*red*) and PyoS5_194-315_ (*grey*) at RT in 20 mM NaCl, 10 mM potassium phosphate buffer pH 7.5 show the PyoS5 domains are helical and folded. (B) CD thermal melts of 0.1 mg/mL PyoS5_1-315_ (*black*) Tm 51.0 ±0.1 °C, PyoS5_1-196_ (*red*) T_m_ 57.5 ±0.0 °C, and PyoS5_194-315_ (*grey*) T_m_ 43.7 ±0.3 °C in 20 mM NaCl, 10 mM potassium phosphate buffer pH 7.5. Two repeats were performed; one is shown as circles with its fit as a line.

**Supplementary Figure S6:**
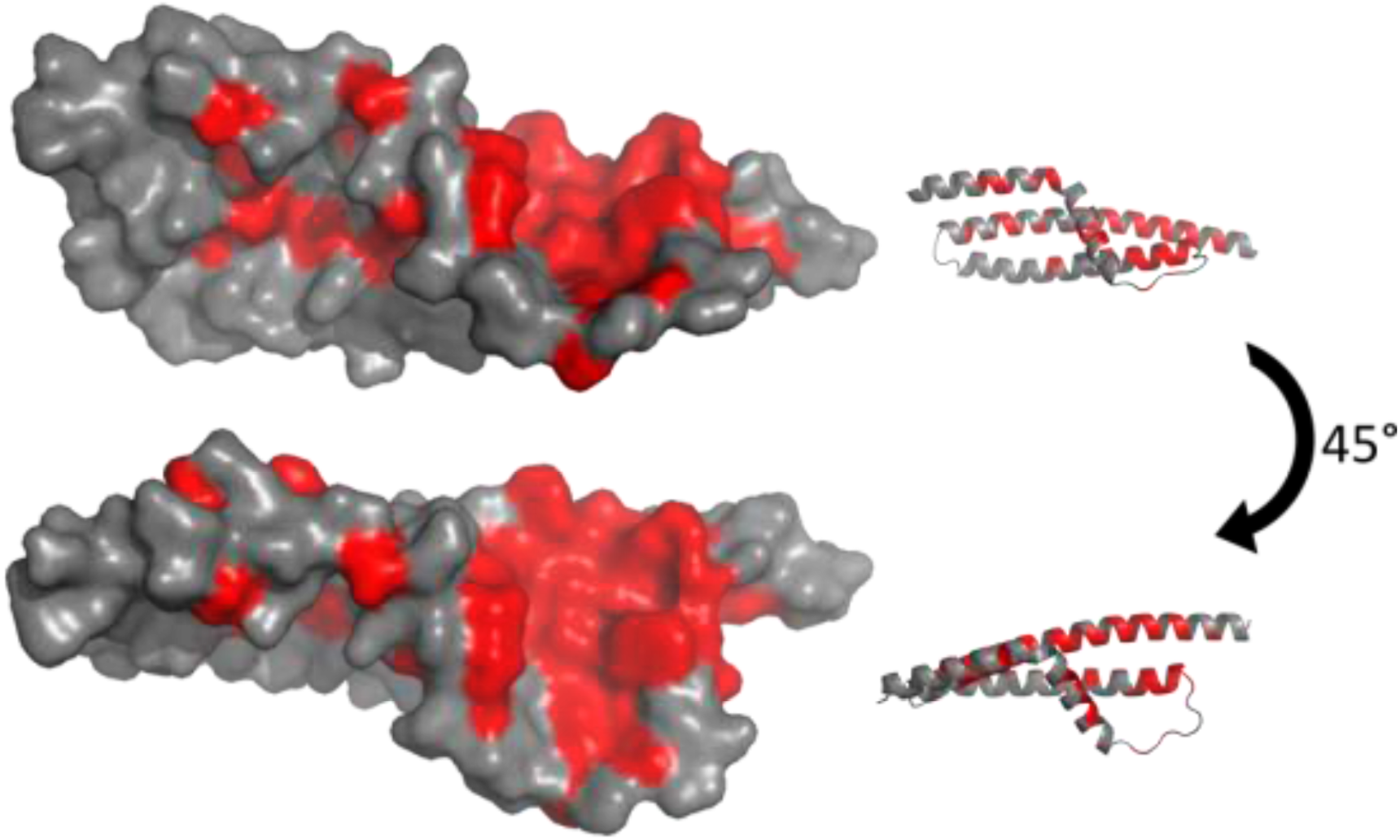
Conserved residues between PyoS2, PyoSD3 and PyoS5 CPA-binding domains. Residues identical in all three CPA-binding domains (*red*) are mapped onto PyoS5 CPA-binding kTHB domain 2 (*grey*). Surface and cartoon representation are shown at two different angles.

**Supplementary Figure S7:**
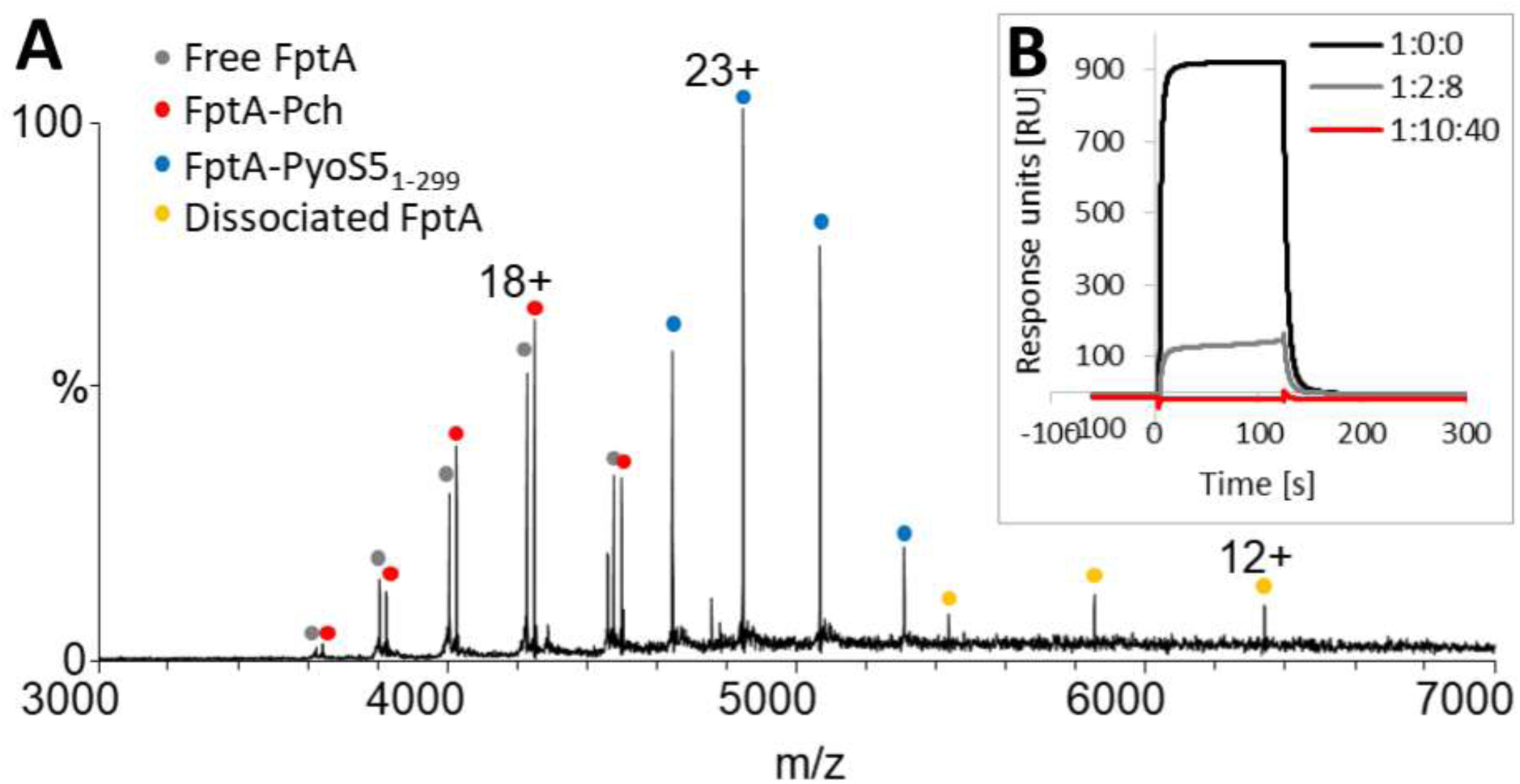
Pyochelin and PyoS5 compete for FptA binding. (A) FptA was incubated with 4-fold excess ferric pyochelin (Pch) for 45 min, then excess Pch was removed by desalting. The native mass spectrum of this FptA sample (19 μM) in the presence of 19 μM PyoS5_1-299_ shows species with masses consistent with free FptA (*grey*), FptA-Pch complex (*red*), FptA-PyoS5_1-299_ complex (*blue*) and dissociated FptA (*yellow*). No complex of FptA, PyoS5_1-299_ and Pch is detected. (B) SPR shows 3.4 μM FptA binding to immobilized PyoS5_1-315_ in the presence of pre-mixed Pch and FeCl_3_ at the following ratios of 1:0:0 (*black*), 1:2:8 (*grey*) and 1:10:40 (*red*) FptA to Pch to FeCl_3,_ respectively. An excess of iron over Pch was chosen to ensure a high percentage of ferric Pch. With a 10-fold excess of Pch no binding of FptA to PyoS5_1-315_ is observed.

**Supplementary Figure S8:**
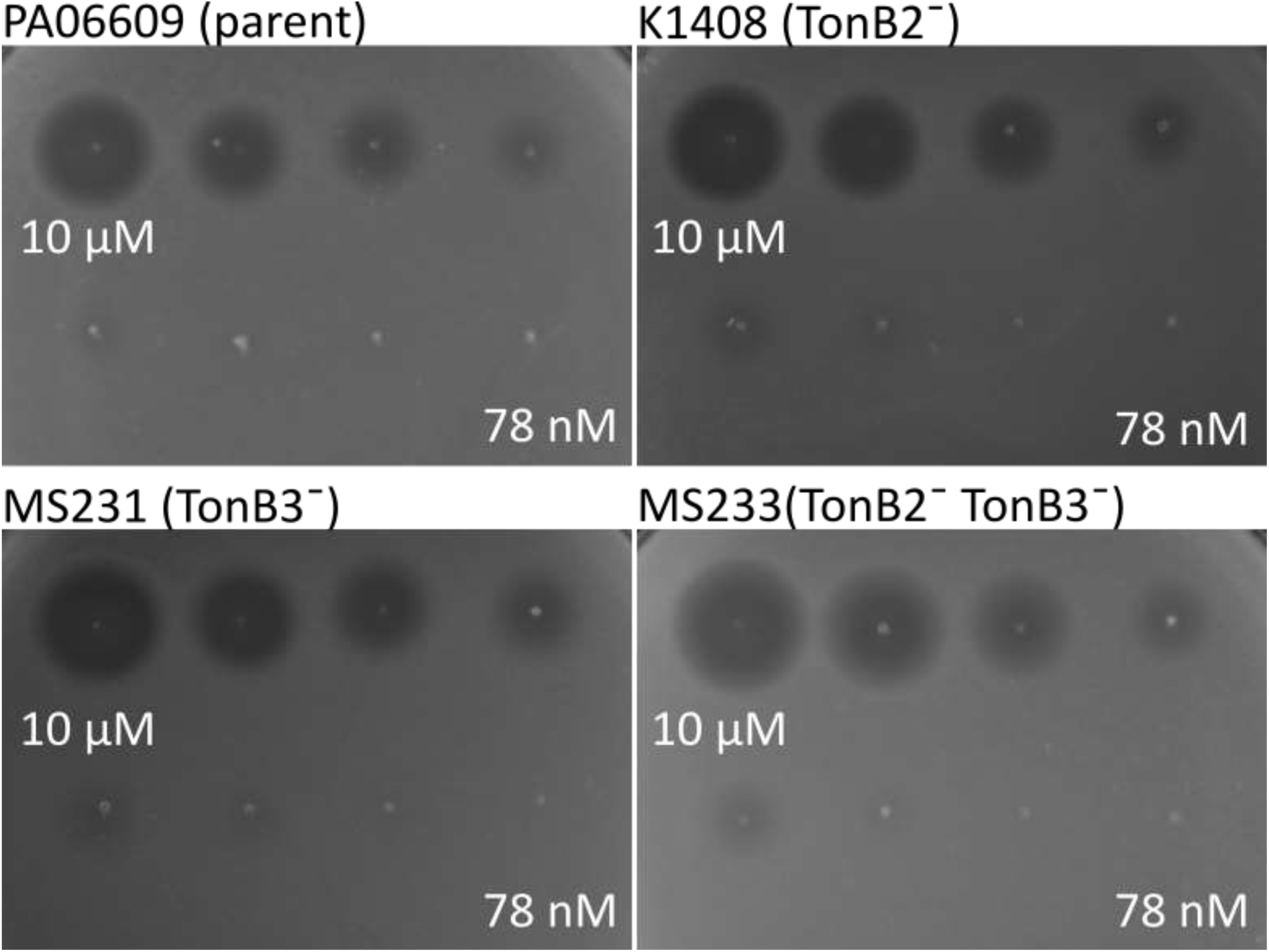
PyoS5ColIa is not TonB2 or TonB3 dependent. Zones of clearance from PyoS5ColIa did not differ between parent strain PA6609, TonB2-deficient strain K1408, TonB3-deficient strain MS231, and TonB2- and TonB3-deficient strain MS233. 3-fold dilution series starting at 10 µM PyoS5ColIa added to strains grown on LB agar.

**Supplementary Figure S9:**
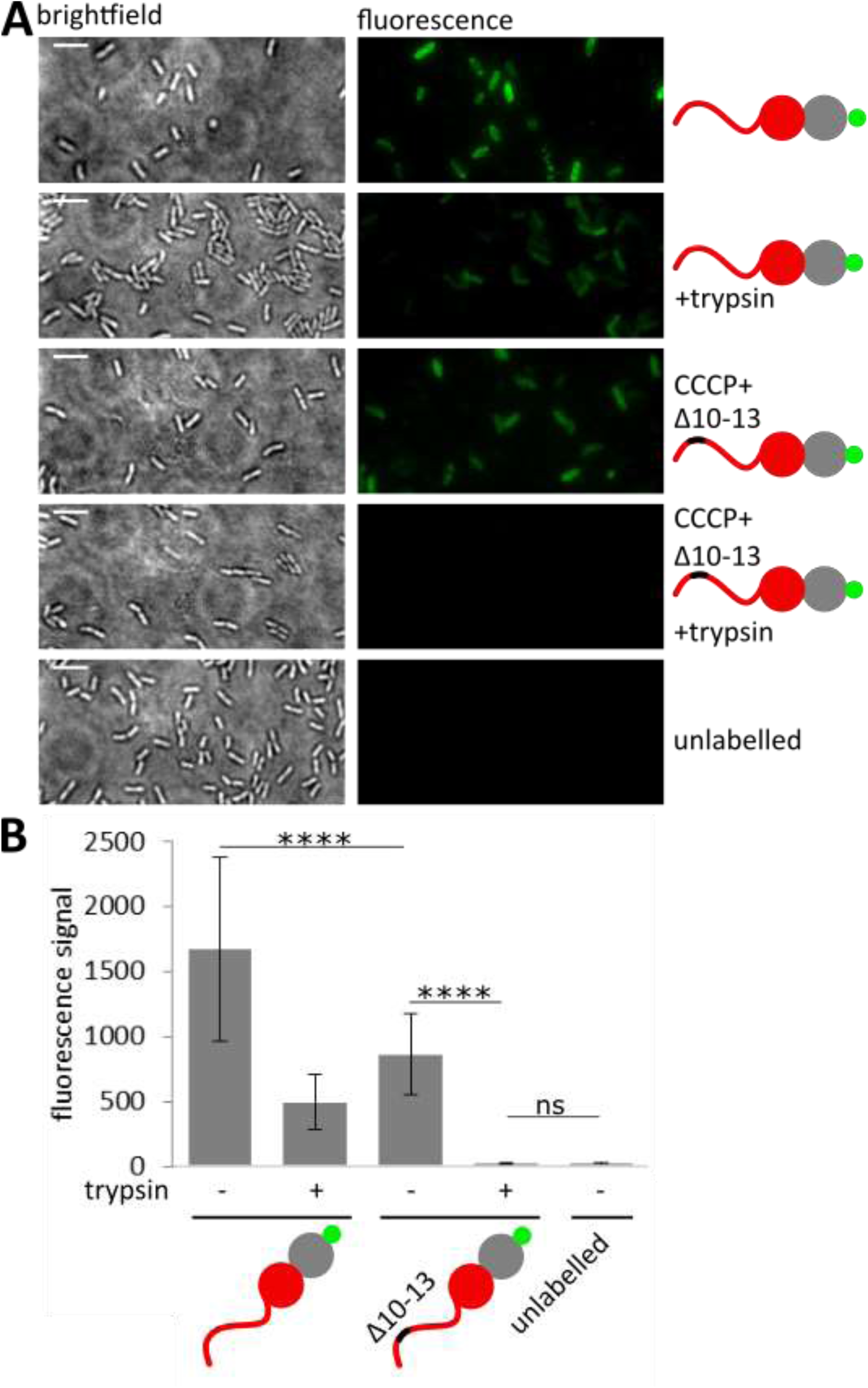
Deletion of the PyoS5 TonB-box prevents translocation. (A) Fluorescent labelling of live *P. aeruginosa* PAO1 using PyoS5_1-315_-AF^488^ and PyoS5_1-315_ Δ10-13-AF^488^ with and without trypsin treatment. Scale bars 5 μm. (B) Quantification of the average cell fluorescence observed under different conditions tested in A. **** indicates a p-value below 0.0001 in Student’s t-test, ns indicates p-value above 0.05.

**Supplementary Figure S10:**
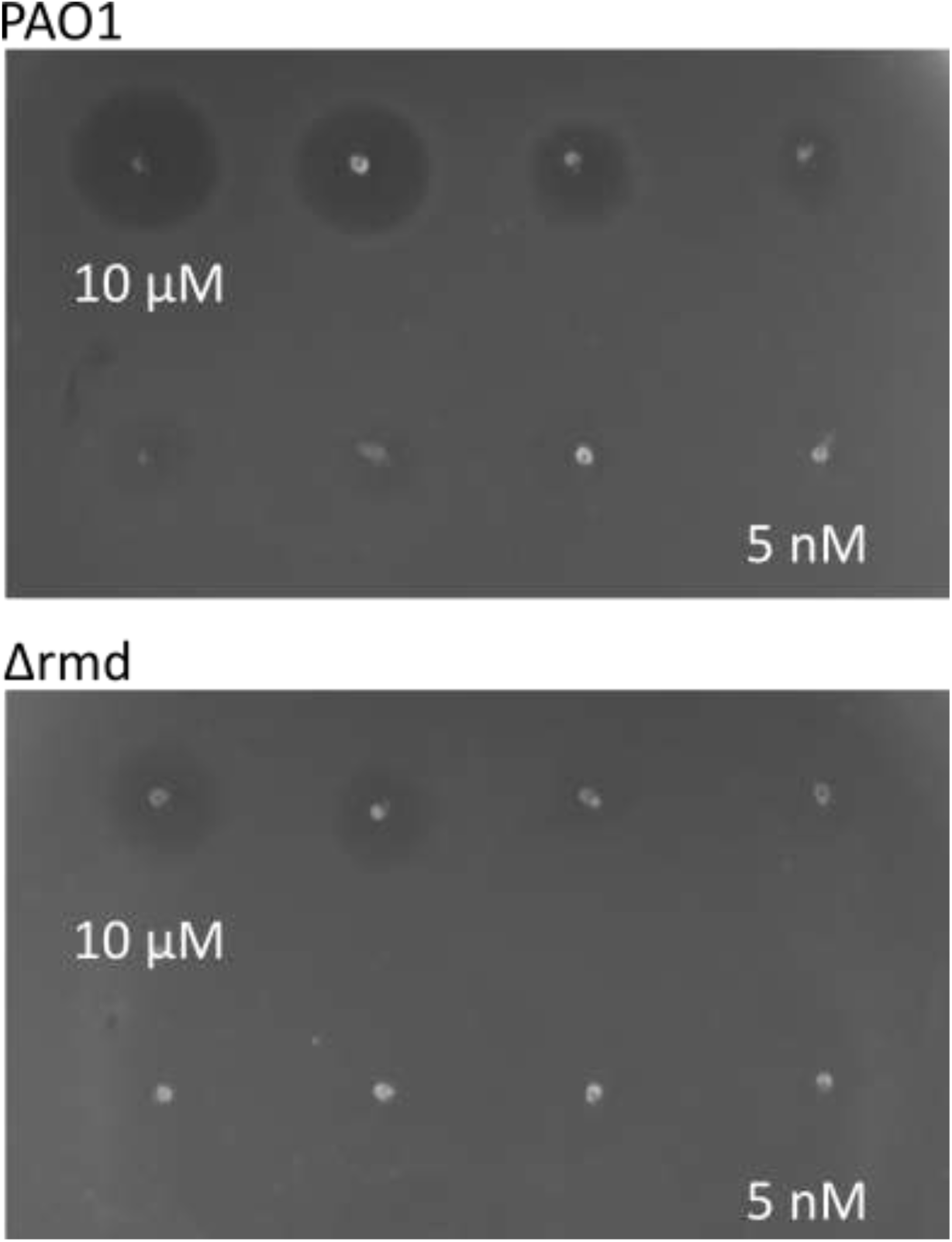
CPA increases susceptibility to PyoS5ColIa. 3-fold serial dilution starting at 10 μM spotted onto the soft agar containing *P. aeruginosa* PAO1 or *P. aeruginosa* PAO1 Δ*rmd* from top left to bottom right. In the CPA-deficient Δ*rmd* strain a 9-fold reduction in susceptibility but not complete resistance is observed.

**Supplementary Figure S11:**
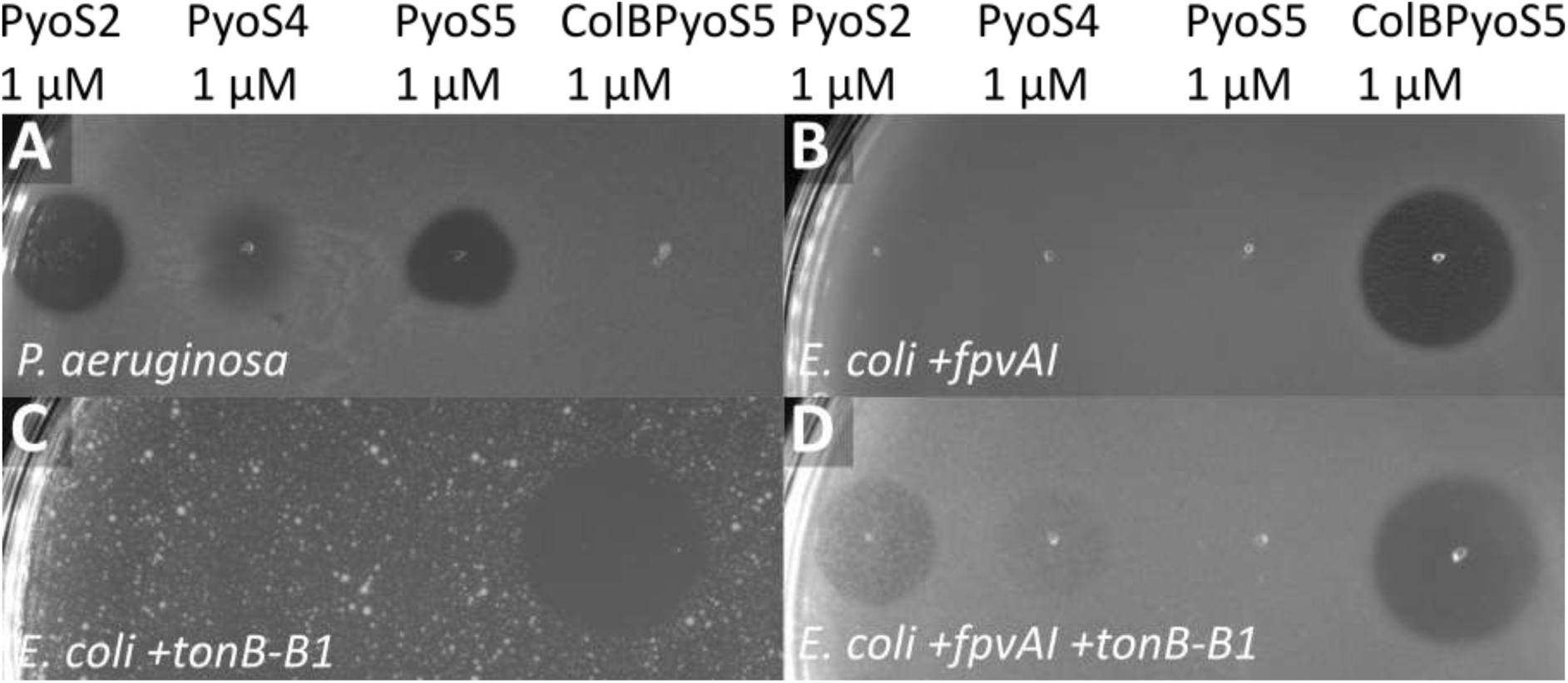
FpvAI and TonB1 constitute the minimal system for PyoS2- and PyoS4-susceptibility. Susceptibility to 1 μM PyoS2, PyoS4, PyoS5 and ColBPyoS5 was assessed for (A) *P. aeruginosa* YHP17, (B) *E. coli* BL21 (DE3) expressing FpvAI, (C) *E. coli* BL21(DE3) expressing TonB-B1, and (D) *E. coli* BL21 (DE3) expressing FpvAI and TonB-B1. Zones of clearance were observed for all *E. coli* conditions tested when ColBPyoS5 was applied (B-D). Similarly, *E. coli* expressing FpvAI and TonB-B1 was susceptible to both PyoS2 and PyoS4, as was the *P. aeruginosa* control (A+D). Zones of clearance for PyoS5 were only observed in *P. aeruginosa*.

**Supplementary Table S1:** X-ray data processing, refinement and validation statistics for PyoS5.

| **Data Processing** |  |
| --- | --- |
| X-ray wavelength | 0.9679 Å |
| Space group | P 1 2<sub>1</sub> 1 |
| Cell dimensions | a = 50.246 Å, b = 52.878 Å, c = 104.807 Å |
| Cell angles | $\alpha = 90^\circ$, $\beta = 95.16^\circ$, $\gamma = 90^\circ$ |
| Resolution | 53.88-2.20 Å (2.27-2.20 Å) |
| Unique reflections | 28285 (2473) |
| Completeness | 98.8 % (99.8 %) |
| Multiplicity | 4.4 (4.6) |
| CC(1/2) | 0.996 (0.728) |
| R<sub>meas</sub> | 0.179 (2.52) |
| **Refinement** |  |
| R<sub>work</sub>/R<sub>free</sub> | 0.225/0.275 |
| Number of protein atoms | 3633 |
| Number of Zn<sup>2+</sup> atoms | 8 |
| Average B-factor | 63.91 Å<sup>2</sup> |
| **Validation** |  |
| RMS bonds | 0.0096 Å |
| RMS angles | 1.05° |
| Ramachandran favoured | 98.49% |
| Ramachandran allowed | 1.51% |
| Ramachandran outliers | 0% |
| MolProbability Clashscore | 2.76 |

Values in parenthesis denote highest resolution shell.

**Supplementary Table S2:** K_d_s of PyoS5 constructs binding to LPS-derived polysaccharides.

| Polysaccharide in the cell with concentration | Titration with concentration | K<sub>d</sub> [nM] | ΔH [kcal/mol] | ΔS [cal/mol/K] | N |
| --- | --- | --- | --- | --- | --- |
| 7 mg/mL<br>*P. aeruginosa*<br>PAO1 LPS-derived polysaccharide containing CPA and OSA | 150 μM PyoS5<sub>1-315</sub> | 612 ±332 | -18 ±6 | -32 ±22 | 0.16 ±0.04 |
| 7 mg/mL<br>*P. aeruginosa*<br>PAO1 LPS-derived polysaccharide containing CPA and OSA | 150 μM PyoS5<sub>1-196</sub> | No binding was observed |  |  |  |
| 7 mg/mL<br>*P. aeruginosa*<br>PAO1 LPS-derived polysaccharide containing CPA and OSA | 150 μM PyoS5<sub>194-315</sub> | 269 ±44 | -14 ±0.3 | -16 ±1 | 0.56 ±0.36 |
| 7 mg/mL<br>*P. aeruginosa*<br>Δrmd LPS-derived polysaccharide containing OSA only | 150 μM PyoS5<sub>194-315</sub> | No binding was observed |  |  |  |

**Supplementary Table S3:**
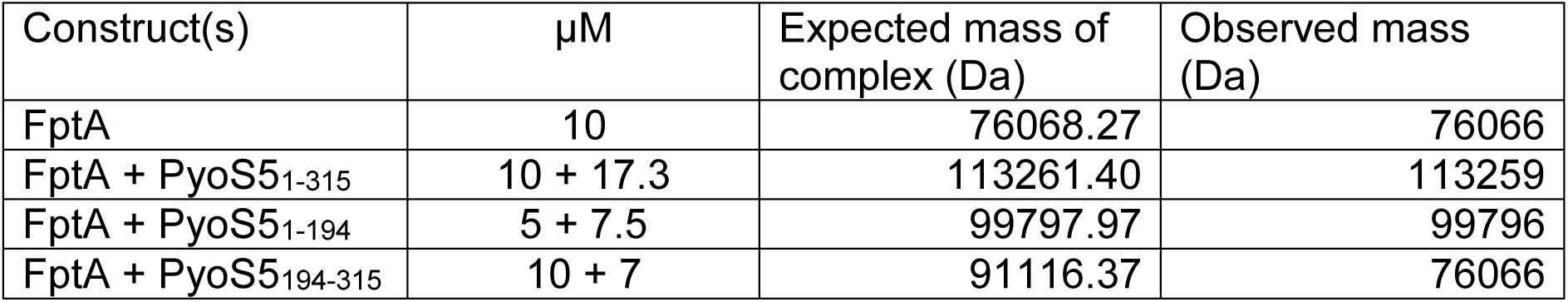
PyoS5_1-196_ binds FptA. Complex formation of FptA with PyoS5 constructs observed by native MS

**Supplementary Table 4:** K_d_s determined by SPR at 25 °C, average of three repeats.

| Ligand immobilized by amine coupling | Amount immobilized [RUs] | Analyte in HBS-OG buffer | $K_d$ |
| --- | --- | --- | --- |
| PyoS5<sub>1-315</sub> | 3464 | FptA | 6.5 ±0.4 μM |
| PyoS5<sub>1-196</sub> | 4153 | FptA | 7.1 ±0.7 μM |
| PyoS5<sub>194-315</sub> | 2088 | FptA | No binding observed |
| PyoS5<sub>1-315</sub> Δ2-39 | 9223 | FptA | 14.7 ±0.4 μM |
| PyoS5<sub>1-315</sub> | 3464 | TonB1 | 241 ±9 nM |
| PyoS5<sub>1-315</sub> Δ10-13 | 5324 | TonB1 | No binding |
| PyoS5<sub>1-315</sub> Δ2-9 | 5352 | TonB1 | 1.57 ±0.09 μM |
| PyoS5<sub>1-315</sub> Δ16-20 | 7727 | TonB1 | 3.12 ±0.05 μM |
| PyoS5<sub>1-315</sub> Δ2-9 | 5324 | FptA | 6.53 ±0.13 μM |
| PyoS5<sub>1-315</sub> Δ10-13 | 5352 | FptA | 10.07 ±0.19 μM |
| PyoS5<sub>1-315</sub> Δ16-30 | 7727 | FptA | 36.2 ±4.9 μM |

